# The structural basis of hyperpromiscuity in a core combinatorial network of Type II toxin-antitoxin and related phage defence systems

**DOI:** 10.1101/2023.03.22.533649

**Authors:** Karin Ernits, Chayan Kumar Saha, Tetiana Brodiazhenko, Bhanu Chouhan, Aditi Shenoy, Julián J. Duque-Pedraza, Veda Bojar, Jose A. Nakamoto, Tatsuaki Kurata, Artyom Egorov, Lena Shyrokova, Marcus J. O. Johansson, Toomas Mets, Aytan Rustamova, Jelisaveta Džigurski, Tanel Tenson, Abel Garcia-Pino, Arne Elofsson, Vasili Hauryliuk, Gemma C. Atkinson

**Author notes:** These authors contributed equally (Karin Ernits, Chayan Kumar Saha). Corresponding authors, Karin Ernits, Vasili Hauryliuk, Gemma C. Atkinson.

## Abstract

Toxin-antitoxin (TA) systems are a large group of small genetic modules found in prokaryotes and their mobile genetic elements. Type II TAs are encoded as bicistronic (two-gene) operons that encode two proteins: a toxin and a neutralising antitoxin. Using our tool NetFlax (standing for Network-FlaGs for toxins and antitoxins) we have performed a large-scale bioinformatic analysis of proteinaceous TAs, revealing interconnected clusters constituting a core network of TA-like gene pairs. To understand the structural basis of toxin neutralisation by antitoxins, we have predicted the structures of 3,419 complexes with AlphaFold2. Together with mutagenesis and functional assays, our structural predictions provide insights into the neutralising mechanism of the hyperpromiscuous Panacea antitoxin domain. In antitoxins composed of standalone Panacea, the domain mediates direct toxin neutralisation, while in multidomain antitoxins the neutralisation is mediated by other domains, such as PAD1, Phd-C and ZFD. We hypothesise that Panacea acts as a sensor that regulates TA activation. We have experimentally validated 16 new NetFlax TA systems. We used functional domain annotations and with metabolic labelling assays to predict their potential mechanisms of toxicity (such as disruption of membrane integrity, inhibition of cell division and abrogation of protein synthesis) as well as biological functions (such as antiphage defence). The interactive version of the NetFlax TA network that includes structural predictions can be accessed at http://netflax.webflags.se/.

**Significance statement:** Toxin-antitoxin systems are enigmatic components of microbial genomes, with their biological functions being a conundrum of debate for decades. Increasingly, TAs are being found to have a role in defence against bacteriophages. By mapping and experimentally validating a core combinatorial network of TA systems and high-throughput prediction of structural interfaces, we uncover the evolutionary scale of TA partner swapping and discover new toxic effectors. We validate the predicted toxin:antitoxin complex interfaces of four TA systems, uncovering the evolutionary malleable mechanism of toxin neutralisation by Panacea-containing PanA antitoxins. We find TAs are evolutionarily related to several other phage defence systems, cementing their role as important molecular components of the arsenal of microbial warfare.

## Introduction

Toxin-antitoxin (TA) systems typically consist of two adjacent, often overlapping genes that encode a toxin whose expression causes growth arrest and a cognate antitoxin that negates the toxic effect [1]. Based on the nature and mode of action of the antitoxin, TA systems are classified into eight types [2]. While TA antitoxins can be either RNA- or protein-based [2], the toxins are near-universally proteinaceous with the exception of rare RNA-based Type IV systems [3, 4]. The most common group of proteinaceous TA pairs is Type II, where the protein antitoxin directly binds to the protein toxin to sequester it into an inert complex [2].

The first TA operon to be discovered, *ccdAB*, was identified serendipitously in 1983 due to its stabilising effect on plasmid maintenance [5], and more TAs were identified *ad hoc* in the following decades [6–10]. The rate of TA discovery has dramatically increased as high-throughput approaches for TA identification have been developed. Systematic experimental discovery of TAs was first achieved using shotgun cloning for identification of toxic ORFs [11]. More recently, novel TAs have been identified through selection for phage immunity phenotypes [12–14]. As the number of sequenced genomes and known TAs have grown, sensitive sequence searching and “guilt by association” – i.e. conserved colocalisation of toxin and antitoxin as a bicistronic operon – have been used for *in silico* discovery of new TA systems [15–23]. Our bioinformatics-driven TA discovery relies on analysis of gene neighbourhood conservation using a sequence search approach that is sensitive enough to find remote similarity even in small, divergent proteins [19, 24, 25].

TA systems are ubiquitous in microbial life, being found encoded on mobile genetic elements and chromosomes of bacteria and archaea, as well as in genomes of temperate phages and prophages [19, 26, 27]. The wide distribution and extreme diversity of TAs has driven the search to discover the biological roles of these systems [28]. The functions that have been put forward over the years can be grouped into three main types. First, addiction or stability modules that support mobile genetic element or genomic region integrity [5, 29]. Second, “emergency brakes” that are activated to stop growth as part of stress responses [30, 31]. Finally, TAs have been discovered to mediate defence against phages via abortive infection [32–34]. The latter function has found substantial recent support. Numerous studies including large-scale exploratory and focused mechanistic approaches have rapidly advanced the field [12, 14, 35–39].

Being frequently horizontally transferred components of accessory genomes, toxin-antitoxin systems have patchy distributions across genomes [15, 40]. It has long been known that Type II toxins and antitoxins have a degree of modularity, in that they can swap partners through evolution [18, 21,40–43]. The hyperpromiscuous antitoxin domain Panacea has provided a recent striking example of how extensive TA partner swapping can be, with <u>P</u>anacea-containing <u>a</u>ntitoxins (PanA) being paired with dozens of different evolutionary and structurally unrelated toxin domains (PanTs) [25]. This discovery suggested that the Panacea domain may have inherent properties that enable it to neutralise multiple unrelated toxins through an unknown mechanism [25]. However, a structural understanding of PanA-mediated neutralisation has been lacking. It is not even clear if the Panacea domain mediates neutralisation directly or merely serves as a platform, with other structural elements providing the inhibitory function. Furthermore, while Panacea’s hyperpromiscuity is remarkable, it is unclear just how much this is paralleled in other antitoxins.

In this study we have systematically explored the TA partner swapping network using NetFlax (standing for <u>Net</u>work-<u>Fla</u>Gs for to<u>x</u>ins and antitoxins), an iterative implementation of our gene neighbourhood analysis tool FlaGs [24], followed by experimental validation and characterisation of novel TA systems. Searching 24,479 representative proteomes from the NCBI RefSeq database, we have identified 3,597 systems within which there are 278 distinct homologous clusters of proteins in 275 distinct combinations of two-gene modules. Validating the approach, we rediscover many classical Type II TA systems such as MqsR/A, VapB/C, RelE/B, Phd/Doc but also reveal novel lineages of unique domain combinations. We have structurally annotated our network of TA-like two-gene architectures through high-throughput prediction of TA complex structures using AlphaFold2 [44] implemented in the FoldDock pipeline [45]. Focusing on the Panacea node of the network, we have validated our structural predictions though extensive mutagenesis. We establish that Panacea is an evolutionally malleable domain that can both inhibit toxins through direct interaction and as serve as a platform for toxin neutralisation by Panacea-associated ZBD (<u>Z</u>n^2+^-<u>b</u>inding <u>d</u>omain) and PAD1 (<u>P</u>anacea-<u>A</u>ssociated <u>D</u>omain 1) domains. The combinatorial network reveals close evolutionary relationships between classical Type II toxin-antitoxin systems and antiphage systems, specifically those that include the AAA+ ATPase and OLD_TOPRIM endonuclease domains such as those seen in PARIS [35], AbiLi [46] the Septu system [47], and ImmA protease-containing systems as seen in RosmerTA systems [13]. We explore the network experimentally through validating 16 new systems in toxicity neutralisation assays and predict their potential mechanisms of toxicity through functional domain annotations and metabolic labelling assays.

## Results

### The NetFlax algorithm reveals a core proteinaceous TA network

To uncover a core framework of the network of TA pairs, we developed the computational tool NetFlax that identifies TA-like gene architectures in an unsupervised manner and generates a toxin-antitoxin domain interaction network. The NetFlax principle is that if one partner gene of a TA system is found in a conserved two-gene neighbourhood with an alternate partner, this is predicted as a new pair, and after “hopping” to this new partner, more partners can be found in the same way. After testing different settings for how conserved a putative TA pair must be to be used as a query in the next hopping step (see **SI Text** and **Figure S1**), we set a requirement that each new pair must be conserved in at least eight representative genomes to be allowed to hop to a new node. As this stringency leads to missing some less well conserved systems, we improved sensitivity though adding a final guilt by association hop for each node, which only required a system to be conserved in two representative genomes.

NetFlax finished hopping after eight hopping steps, converging on a final network, having reached dead-ends for all the network lineages (**Fig. 1**). We initially identified 79 clusters conserved in a minimum of eight genomes. These we call D nodes (standing for central <u>D</u>omain nodes). After the subsequent less strict node analysis allowing conservation in two genomes with no onward hopping, we discovered 234 additional nodes, which we refer to as M nodes, for <u>M</u>ininodes. In total we identified 314 nodes. Toxin/antitoxin assignments are made by virtue of their lineages from the original Panacea antitoxin domain, assuming that the hopping goes from antitoxin to toxin to antitoxin etc. This assumption seems to work well on the whole; in the classical Type II part of the network, our annotation of whether the cluster is a toxin or antitoxin domain matches that in the TADB database [48], and domain annotations (**Dataset S1**). However, we cannot be sure that our annotations hold true for the termini of the network. One extended lineage of three D nodes leading from the Rosmer/ImmA zone (D41, D95, D127 and D132 associated with 32 combined M nodes) became particularly complicated with node domain fusions, making our ability to predict toxins and antitoxins particularly troublesome. Therefore, we decided on balance to “prune” this lineage from the core TA network of **Fig. 1** (however, these lineages and their data are still available in the unpruned interactive network http://netflaxunpruned.webflags.se/ and **Dataset S1**).

**Figure 1.**
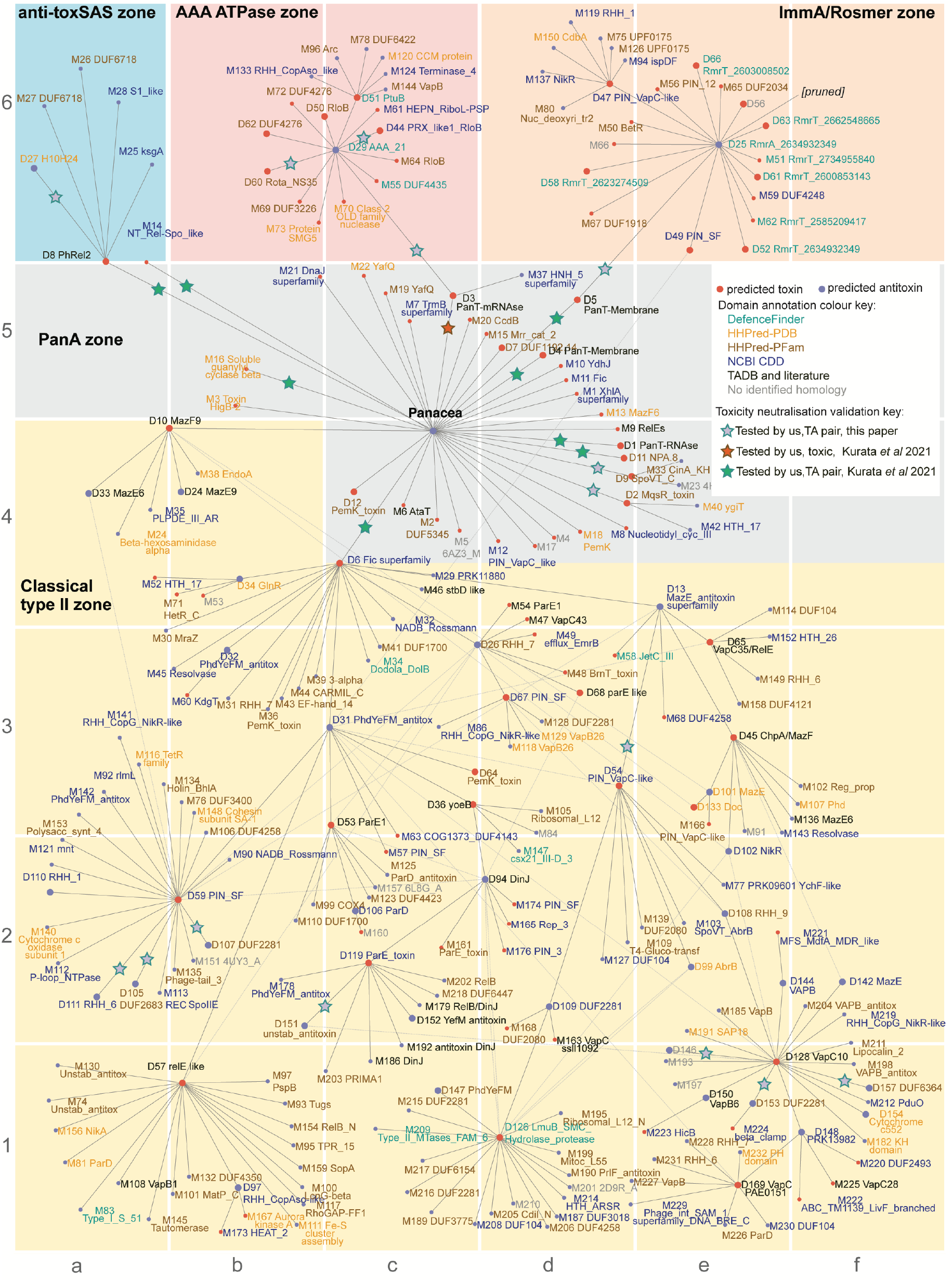
The core proteinaceous TA network. The core network of NetFlax-predicted toxin and antitoxin-like domains across microbial life. The starting input was the Panacea domain. Blue circles are predicted antitoxins and red are predicted toxins. Yellow stars show groups that contain previously verified toxins and antitoxins (**Dataset S1**) [48]. Orange and green stars show, respectively toxins and TAs that have been previously validated, including by us [19, 49]. Predicted toxin and antitoxin domains are annotated based on sequence homology searches (see text for references).

Our final core TA network (**Fig. 1**, http://netflax.webflags.se/) represents the most conserved systems of the 24,474 representative predicted proteomes considered. The network comprises 278 nodes, of which 107 are predicted to be toxins, and 171 are predicted to be antitoxins. These fall into 275 distinct toxin-antitoxin node combinations. It is useful to roughly divide the network into five topological zones: i) the Panacea domain-containing systems at the core of the network, including six systems experimentally validated in our previous analyses (**Dataset S1**) [49], ii) the anti-toxSAS zone containing toxins related to RelA/SpoT alarmone synthetases that likely modify tRNA (nodes D8 and M14), plus their antitoxins, iii) a zone containing a hub AAA ATPase antitoxin domain (D29, **Fig. 1** coordinates c6), iv) a zone containing a hub ImmA protease antitoxin domain (D25, **Fig. 1** coordinates e6) and v) the largest zone of the network where many classical Type II TA systems are found, with considerable interconnections among nodes indicating considerable partner swapping. Panacea-containing systems are the largest group of our TAs (**Fig. 2**), closely followed by D29 (AAA ATPase-like) and D31 (Phdrelated antitoxins).

**Figure 2.**
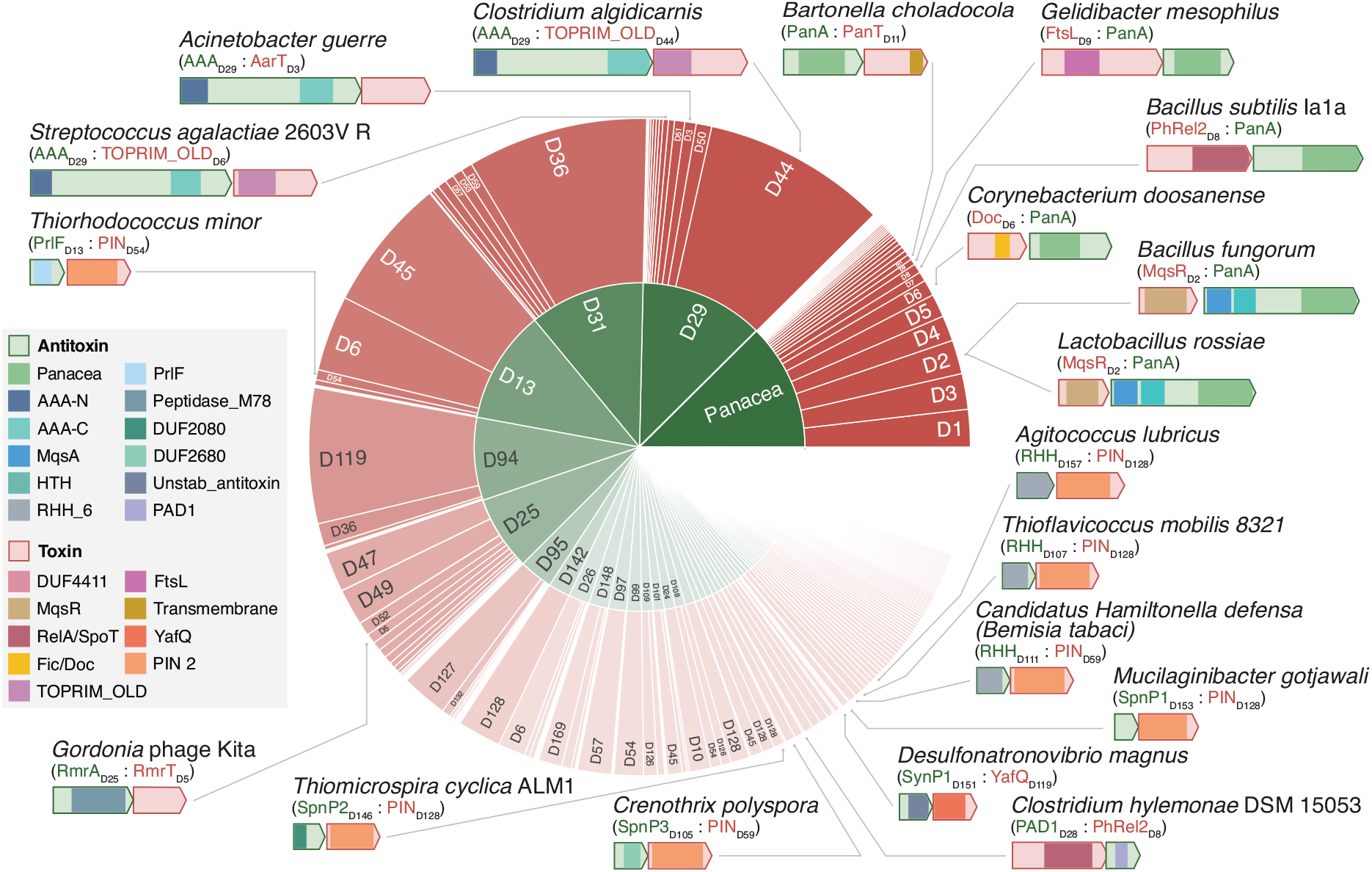
The diversity of systems in the core NetFlax TA network. The size of the sections represents the number of proteins in each cluster (node) of **Fig. 1**. The block arrows represent open reading frames, drawn to scale. Coloured boxes within the arrows indicate domains, coloured according to the legend in the lower right.

### Phage defence systems are widespread in the NetFlax TA network

For each node in the NetFlax network, we made functional predictions for a protein representative by searching with domain models including those from DefenceFinder, a database of phage defence systems [43]. The latter search revealed that the AAA ATPase and ImmA zones are particularly enriched in phage defence systems. The AAA ATPase domain of node D29 (**Fig. 1** coordinates c6) is found in a number of defence systems, such as AriA of the PARIS system [35], GajA of the Gabija system [50] and PtuA of the Septu system [47]. GajA is a sequence-specific ATP-dependent DNA endonuclease that is inhibited by dNTP and NTP nucleotides [50]. It is a two-domain protein, with an N-terminal AAA domain and a C-terminal TOPRIM (<u>to</u>poisomerase-<u>prim</u>ase) endonuclease domain, closely related to OLD (<u>O</u>vercome <u>L</u>ysogenisation <u>D</u>efect) families [50]. In our network, we see the latter domain can be associated with the AAA domain as a separate protein (node M70, coordinates c6). This is the same two-gene architecture as seen in AriAB of the PARIS system [35]. the Among the other nodes linked to D29 is D44, homologous to RloB. The RloB protein family has been observed in Type I restriction-modification operons [51], and the AbiLii protein, which is part of a plasmid-encoded phage abortive infection mechanism [46]. HHPred [52] indicates the RloB domain is also related to OLD_TOPRIM domains. The node D51 is homologous to the HNH nuclease domain, as seen in Septu protein PtuB (**Dataset S1**). Thus D29 and its cognate D51 together constitutes a similar two-domain PtuAB Septu system architecture previously identified in *B. thuringiensis* [47]. The presence of the HEPN nuclease domain in node M61 indicates a general tendency for nuclease domains to be associated with AAA ATPase domains.

The ImmA protease domain which NetFlax predicts as an antitoxin (D25, **Fig. 1** coordinates e6) was recently confirmed as such in the diverse phage defence RosmerTA systems, where different toxins (RmrTs) are paired with the protease domain-containing antitoxin (RmrA) [12, 13, 53] (**Fig. 1**). The ImmA domain is named after the protein encoded on a conjugative transposon of *Bacillus subtilis* [54]. ImmA is an anti-repressor that cleaves an HTH (<u>H</u>elix-<u>T</u>urn-<u>H</u>elix) domain-containing repressor (ImmR) to allow the expression of an integrase [54]. Indeed, proteins within the D25 node often possesses a small N-terminal HTH domain in addition to the protease domain, suggesting a similar HTH cleavage mechanism of regulation in Rosmer-like TAs. Again, we see the involvement of nucleases in our predicted systems, this time in an association of D25 with PIN-like domains, the most diverse and ubiquitous nuclease superfamily, often seen as toxin components of TAs [55, 56].

Phage defence domains appear in other parts of the NetFlax network. In addition to the AAA and ImmA/Rosmer zones that are clearly phage defence related, we have recently discovered that toxSASs can protect against phages [39]. Additionally, the identification of DefenceFinder domains in the Classical Type II zone, along with evidence for classical TA domains in phage defence [57, 58] shows how intricately TAs in general are associated with phage defence.

### NetFlax TAs are found across the prokaryotic tree of life, and in tailed bacteriophages

NetFlax-predicted TAs are found in all major phyla of bacteria and archaea (**Fig. S2**, **Dataset S1**). Most NetFlax TAs were found in Pseudomonadota (formally known as Proteobacteria) – particularly Gammaproteobacteria, reflecting the bias of RefSeq towards these taxa. The pseudomonad *Thiobaca trueperi* has the most NetFlax-predicted TAs (eight). NetFlax TAs are also found across the archaeal tree of life, with representatives in the phyla Euryarchaeaota, Crenarchaeota, Thaumarchaeota, Candidatus Thermoplasmota and Candidatus Koracheota. Within viruses, TAs were only predicted in Uroviricota (tailed bacteriophages). Our discovery of 13 NetFlax TAs in phages (**Dataset S1)** is likely a significant underestimate as many bacteria-encoded systems are likely resident on prophages integrated into the bacterial chromosome.

### AlphaFold2 confidently predicts the structure of binary TA complexes

TAs are excellent targets for modern deep-learning structural prediction methods – not just of single proteins – but of complexes. This is because Type II systems necessarily form tight complexes to keep the toxin in check, with a co-evolutionary signal in the interface region [59]. AlphaFold2 is a particularly powerful structural prediction method that takes advantage of the co-evolution of residues for prediction [44]. We have run AlphaFold2 on a high throughput basis with the FoldDock pipeline [45] to predict the structure of all 3,597 protein pairs (3,277 after pruning). To keep predictions computationally feasible, we predict binary TA dimers, not higher order oligomers, rationalising that even in larger complexes, there must be an interface between the toxin and antitoxin. The reliability of the structures of the complexes is assessed using the pDockq score, which takes into account the number of interface contacts, and the plDDT reliability scores from AlphaFold2 for those regions. All structures and their scores are available on the interactive network (http://netflax.webflags.se/) Recapitulation of TA folds previously solved with X-ray crystallography indicates these predictions are reliable (**Fig. S3**). We determined the distribution of model confidence (pDockQ scores) of TA pairs compared to random pairs. The TA predictions are much better than random predictions for the same set and roughly 50-60% of the complexes are well modelled (**Fig. S4**).

### Recurrent structural folds appear across the NetFlax network

The NetFlax algorithm includes a cross-checking step for the identified toxin and antitoxin clusters, to determine whether the potential cluster is unique or similar to any cluster discovered during the previous hopping rounds (see **Fig. S1B**). Despite this, we found multiple clusters in the classical Type II zone of the network with similar domain annotations, suggesting they may be homologous domains that are not clustered together. For example, the Phd, ParE and PIN domain appear multiple times in the network (**Fig. 1**). Sequence alignments show that these unclustered but related nodes are clearly distinct in terms of sequence (including insertions and deletions, **Fig. S5**), but that they are similar enough that they have the same fold. To systematically address this, we clustered all our predicted structures and annotated our network to show nodes that have the same fold (**Fig. S6**). We found many of our predicted TAs can be clustered into 18 distinct folds, the most common being ATPase, MazF/PemK, PIN, RelE/ParE, Phd/YefM, and a common fold of Rosmer toxins in the DefenseFinder database [43]. Structural alignments of the most common folds are shown in **Fig. S7**. This supports previous observations that toxins and antitoxins that are diverse at the sequence level can have the same structural fold [60]. A similar conservation of domains is also seen in phage defence systems [61].

### PanA-containing TAs: the roles of individual antitoxin domains in toxin neutralisation

We focused on four previously experimentally validated PanTA TAs: *Bartonella choladocola* (previously *Bartonella apis*) PanT_D11_:PanA, *Corynebacterium doosanense* Doc_D6_:PanA, *Bacillus subtilis* Ia1a toxSAS PhRel2_D8_:PanA and *Bacillus fungorum* MqsR_D2_:PanA. The subscript D number refers to the node in **Fig. 1.** All NCBI protein accession numbers of TAs characterised in this paper are shown in **Table S1**. In all of the systems, the Panacea domain has the same compact architecture comprised of α-helices α1-α7 and ß-strands β1 and β2 (**Fig. 3AB**). Despite these PanTAs having dramatically different toxins, these four TA structures are predicted with confidence (pDockQ sores from 0.68 to 0.71). As selected PanTA systems differ in their antitoxin architecture (**Fig. 3C-F**, and see below), mutational analysis of the set allows us to interrogate the function of Panacea antitoxin in PanAs: does it mediate the toxin neutralisation directly or is it merely a ubiquitous accessory domain that has other functions, such as transcriptional regulation of the TA locus or sensing toxin-activating stimuli?

**Figure 3.**
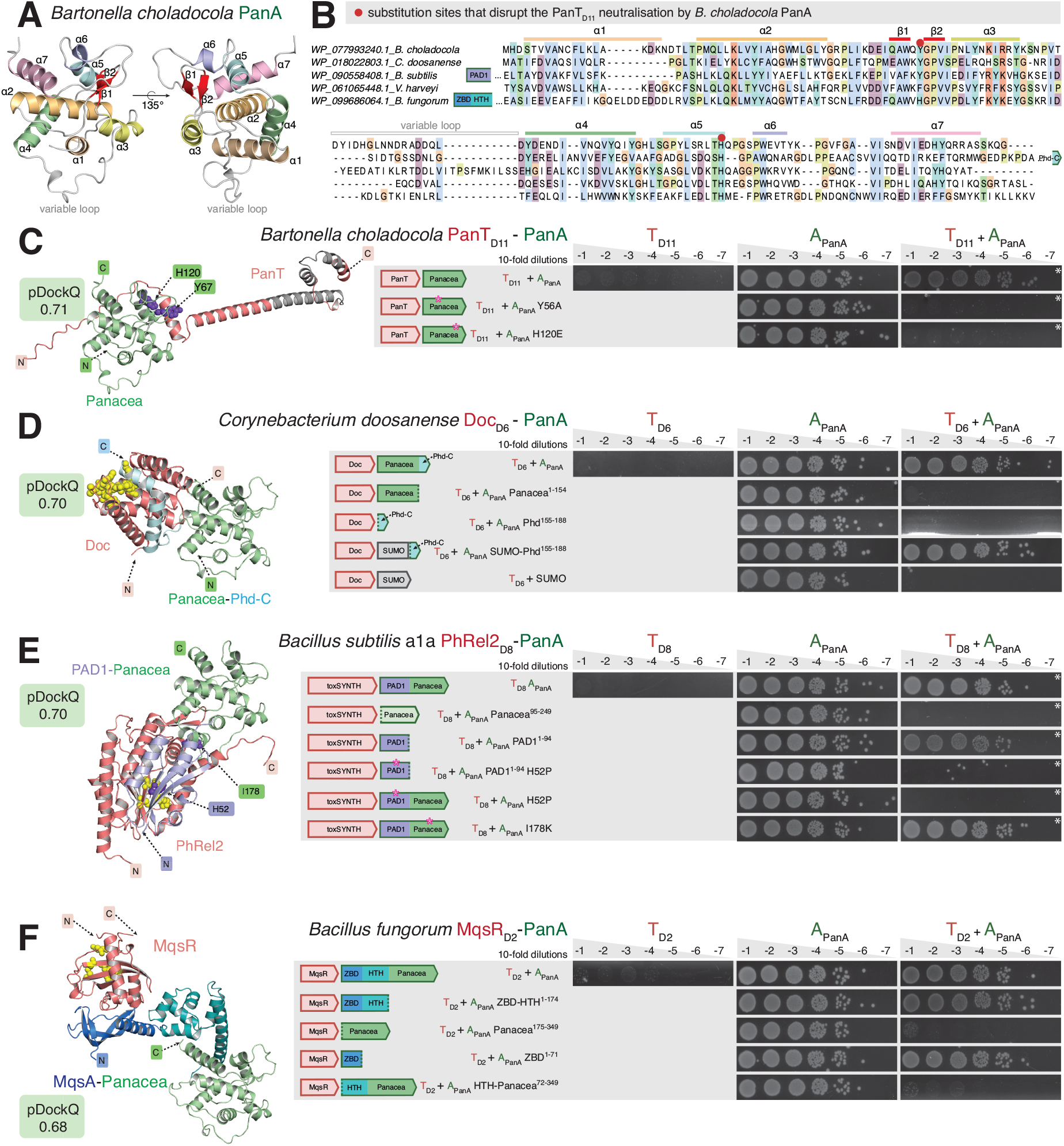
PanA antitoxins employ neutralise PanT toxins either via Panacea domain directly or via additional N-terminal domains: Phd-C, PAD1 and MqsA. (**A**) AlphaFold-generated structural model of *Bartonella choladocola* PanA. Helixes and beta-sheets are labelled as per Kurata and colleagues [25]. (**B**) Alignment of Panacea domains from representative PanA antitoxins. (**C-D**) Mutational probing of AlphaFold-generated PanTA structural models. In toxicity neutralisation assays overnight cultures of *E. coli* strains transformed with pBAD33 and pMG25 vectors or derivatives expressing putative *panT* toxins and *panA* antitoxins, correspondingly, were adjusted to OD_600_ 1.0, serially diluted from 10^1^- to 10^8^-fold and spotted on LB medium supplemented with appropriate antibiotics and inducers (0.2% arabinose for *panT* induction and 0.05 or 0.5 (*) mM IPTG for *panA* induction). Predicted transmembrane domains are shown in grey and the active center of the toxin is highlighted in yellow. Introduced mutations on PanA antitoxin are shown in purple. Protein accessions are in **Table S1.**

The structure of *B. choladocola* PanT_D11_:PanA suggests that Panacea can, indeed, directly neutralise the toxin (**Fig. 3C**). In this case Panacea is predicted to form a contact with the N-terminal unstructured region as well as a short a-helix that precedes the PanT_D11_ predicted transmembrane region [25]. Substitutions Y56A (N-terminally adjacent to β2) and H120E (C-terminal end of α5) that were designed to disrupt this interface do, indeed, render *B. choladocola* PanA unable to neutralise the toxin, thus supporting the structural model. Both of these substituted residues are located in the conserved structural core of the Panacea domain.

In the case of *C. doosanense* PanTA, the two additional C-terminal helices decorating the Panacea core of the PanA antitoxin are predicted to make extensive contact with Doc_D6_ toxin (**Fig. 3D**). The globular Panacea domain itself is not predicted to be involved in neutralisation. These two C-terminal helices are structurally analogous to those found in the C-terminal extension of the *E. coli* Phd antitoxin that inhibits the Doc toxin [62]. Therefore, we refer to this element of *C. doosanense* PanA as the Phd-C domain. *E. coli* Doc is a kinase that phosphorylates EF-Tu to abrogate the cellular protein synthesis [63]; *C. doosanense* Doc_D6_ similarly targets translation [25], and the active site residues are conserved amongst the two proteins (**Fig. 3D**). The Phd-C domain of PanA directly interacts with the active site of the toxin. Truncation of the Phd-C domain renders *C. doosanense* PanA unable to neutralize the toxin (**Fig. 3D**). Expression of the isolated Phd-C domain does not neutralise the toxin, which could be due to the intrinsic instability of the element. To test this hypothesis, we fused Phd-C with a stabilising N-terminal SUMO tag, and as predicted, the resulting construct can readily neutralise Doc_D6_, despite lacking the Panacea domain. No neutralisation was observed in the control experiment with SUMO alone. Collectively, these results suggest that Panacea can serve as an accessory domain, with neutralisation being mediated by a dedicated separate domain.

Next, we characterised the *B. subtilis* Ia1a PhRel2_D8_:PanA system. In our previous analysis of the Panacea domain distribution, we discovered a novel domain that we named the PAD1 domain, standing for <u>P</u>anacea-<u>a</u>ssociated <u>d</u>omain 1 [25]. Apart from two strains of Ruminococcaceae where the putative toxin is an ATPase, PAD1-Panacea multidomain PanA antitoxins are only found paired with toxSASs such as PhRel2_D8_, where it is the most widespread antitoxin for this kind of toxin in the NetFlax network. The second most widespread is NetFlax domain D27 (**Fig. 1**). Remarkably, structural alignment of the toxSAS:D27 of *Clostridium hylemonae* DSM 15053 with toxSAS:PAD1-PanA of *Bacillus subtilis* Ia1a, fused TA CapRel [39] showed that D27, PAD1 and pseudo-ZBD have the same fold, and share the same interface with the toxSAS toxin (**Fig. S8**). Importantly, it is PAD1 that forms most of the contacts with the PhRel2_D8_ toxin (**Fig. 3E**). Strikingly, *B. subtilis* Ia1a PAD1 domain alone – with Panacea removed – can neutralise the toxin, thus directly supporting the structural prediction. Furthermore, H52P substitutions that are predicted to break the PAD1:PhRel2_D8_ interface completely abrogate the neutralisation, both in the context of full-length PanA and isolated PAD1. Finally, the I178K substitution located on the Panacea: PhRel2_D8_ interface did not affect the efficiency of neutralisation. To further support the role of PAD1 as a dedicated toxin-neutralising domain, we have, via toxicity neutralisation assays, validated the *Clostridium hylemonae* DSM 15053 TA system comprised of PhRel2_D8_ and the PAD1_D27_ antitoxin (**Fig. S9**). As the *C. hylemonae* antitoxin naturally lacks the Panacea domain, this observation further supports PAD1 being a directly neutralising antitoxin element.

Finally, we have dissected the *B. fungorum* MqsR_D2_:PanA system (**Fig. 3F**). MqsR is an RNase [60, 64] that is neutralised by antitoxin MqsA comprised of an N-terminal <u>Z</u>n^2+^-<u>b</u>inding <u>d</u>omain (ZBD) and C-terminal HTH [60]. While the ZBD interacts with MqsR and inhibits it without directly interacting with the RNase active site, the HTH region dimerises and acts as a transcriptional auto-regulator of the *mqsRA* operon. *B. fungorum* PanA also contains the ZBD-HTH domain composition characteristic of the MqsA antitoxin, with the Panacea domain added to the C-terminus. Our truncation analysis shows that, indeed, also in the case of *B. fungorum* PanTA, the ZBD directly mediates neutralisation of MqsR_D2_. While both isolated ZBD and ZBD-HTH segments efficiently neutralise MqsR_D2_, neither Panacea alone nor HTH-Panacea are sufficient for neutralisation.

Collectively, our results demonstrate that while Panacea can act as a direct toxin neutraliser, it is unlikely to act as such in PanA antitoxins that contain additional dedicated neutralisation domains such as PAD1 or ZBD. Furthermore, the example of *B. fungorum* MqsR_D2_:PanA system suggests that Panacea probably does not act as a transcriptional auto-regulator of *panAT* operons either, as *B. fungorum* PanA contains a dedicated DNA-binding regulatory domain, HTH (as do many other PanAs, [25]). Therefore, we favour the hypothesis that the Panacea domain acts as a sensor responding to – as yet unknown – TA-activating cues.

### Experimental exploration of the NetFlax network

Finally, we explored our NetFlax TA network experimentally. We focused on novel TA pairs, i.e. the ones containing either i) novel toxin or/and antitoxin domains or ii) novel combinations of previously validated toxins and antitoxins. We have validated 15 TA pairs in toxicity neutralisation assays (**Fig. 1**). For 13 of them, we performed metabolic labelling assays with ^35^S methionine (a proxy for inhibition of translation), or ^3^H uridine (a proxy for inhibition of transcription) or ^3^H thymidine (a proxy for inhibition for replication) (**Fig. 1**).

The TA system from *Gelidibacter mesophilus* is comprised of a relatively large (281 aa) toxin T_D9_ (toxFtsL_D9_) paired with a PanA antitoxin (**Fig. 4A**). Similarly to *B. choladocola* PanT_D11_:PanA, *G. mesophilus* PanA is comprised of a stand-alone Panacea domain that directly neutralises the toxin. While the toxin is clearly very efficient in abrogating the formation of bacterial colonies on solid LB plates, induction in liquid culture does not result in rapid growth inhibition nor do we see any dramatic effects in metabolic labelling assays (**Fig. 4B**). The toxin contains an α-helical FtsL domain which is predicted to dimerise and be localised to the cell membrane (**Fig. 4C,D**). FtsL is an essential component of bacterial dividosome which forms a trimeric complex with FtsB and FtsQ via leucine zipper-like (LZ) motifs [65]. Given the partial homology with FtsL, we propose naming the *G. mesophilus* T_D9_ toxin toxFtsLD9. It is tempting to speculate that *G. mesophilus* T_D9_ could act by directly interfering with cell division. Experiments with liquid cultures of *E. coli* expressing *G. mesophilus* toxFtsLD lend support to this hypothesis: after an hour of uninhibited growth, the OD_600_ increase stops and then the culture collapses, suggestive of cell lysis (**Fig. 4E**).

**Figure 4.**
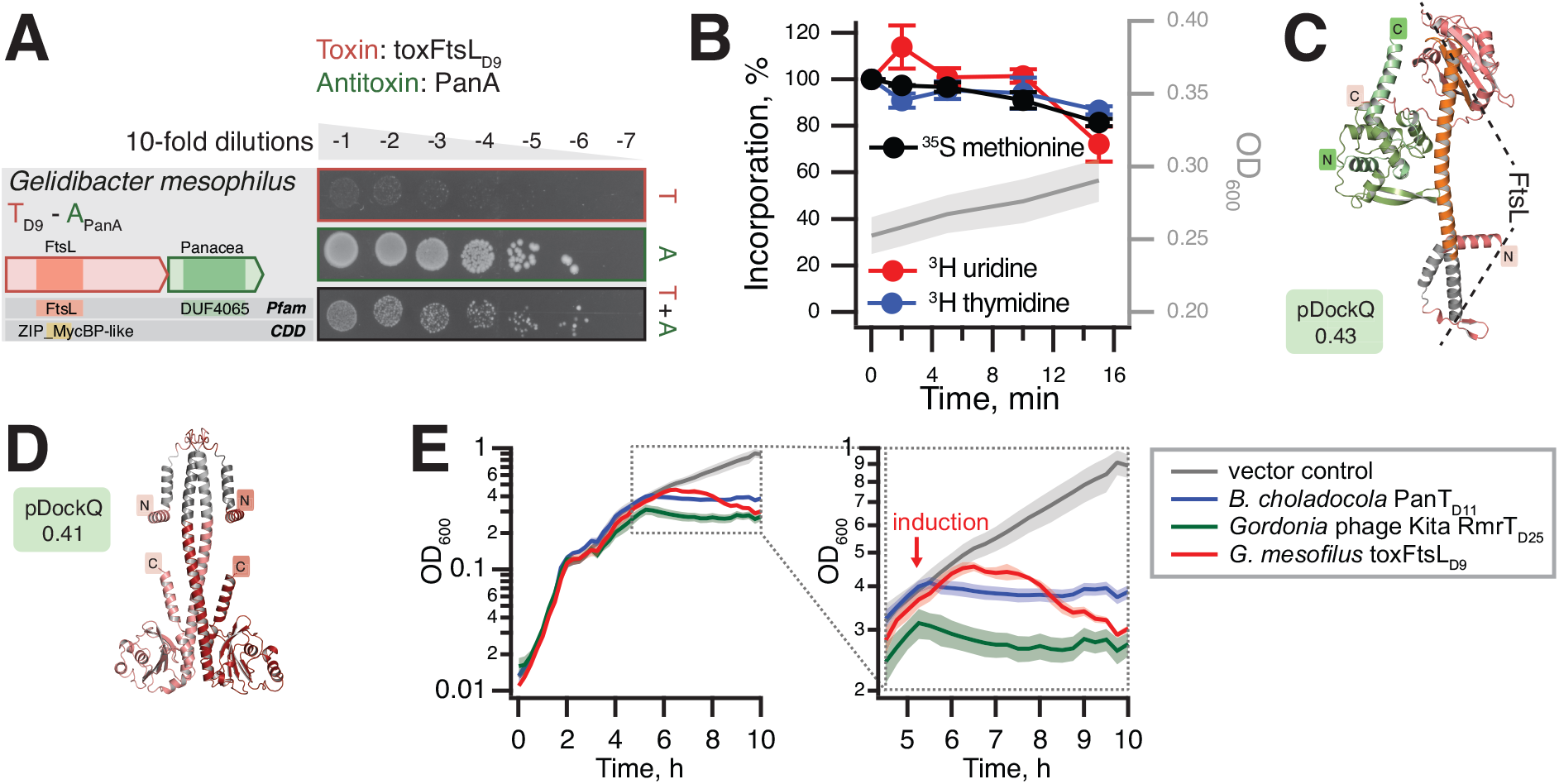
*G. mesophilus* toxFtsL is a slow-acting PanT toxin with partial homology with the FtsL component of bacterial dividosome. (**A**) Validation of the *G. mesophilus* toxFtsL:PanA TA through toxicity neutralisation assay. (**B**) Metabolic labelling assays with wild-type *E. coli* BW25113 expressing *G. mesophilus* toxFtsL. (**C**) AlphaFold-generated PanTA structural model of *G. mesophilus* toxFtsL:PanA TA pair and (**D**) toxFtsL_D9_ dimer model. Predicted transmembrane helical regions are shown in grey, and α-helical FtsL-like region is highlighted with a dotted line. (**E**) Delayed growth inhibition and cell lysis by *G. mesophilus* toxFtsL. Growth assays of *E. coli* cells expressing *G. mesophilus* toxFtsL, *B. cholaadocola* PanT_D11_ or *Gordonia* phage Kita RmrT_D25_ as well as a vector control strain harbouring pBAD33 and pMG25 in MOPS liquid medium supplemented with 0.5% glycerol and 25 μg/ml each 20 amino acids. Expression of toxins was induced with 0.2% arabinose at OD_600_ of around 0.4.

The TA system from *Gordonia* phage Kita is a new member of the RosmerTA family [12, 13, 53] (**Fig. 5A**). The RmrA T_D25_ protease antitoxin is paired with a D_5_ toxin which has no detectable similarity to other protein families. Metabolic labelling assays show rapid and dramatic abrogation of translation, transcription and replication upon expression of the Kita phage D_5_ toxin (**Fig. 5B**). The toxin is not fully neutralised by the antitoxin and the structure of the TA complex can not be reliably predicted by AlphaFold (pDockQ score of 0.05) (**Fig. 5C**). The C-terminal region of the toxin is predicted to be localised in cellular membrane (**Fig. 65C**). In liquid culture experiments expression of RmrTD5 immediatly abrogates bacterial growth without causing a consequent collapse of OD_600_; the toxin is likely to share the mechanism of toxicity with membrane-depolarising *B. choladocola* PanT_D11_ [25] that we used as a control (**Fig. 4E**). Following the nomenclature for RosmerTA toxins [12, 13, 53], we renamed the Kita phage toxin RmrT_D5_. As other RosmerTA systems have been shown to be phage defence systems [12, 13, 53], it is likely that the Kita phage RmrTA has a similar function.

**Figure 5.**
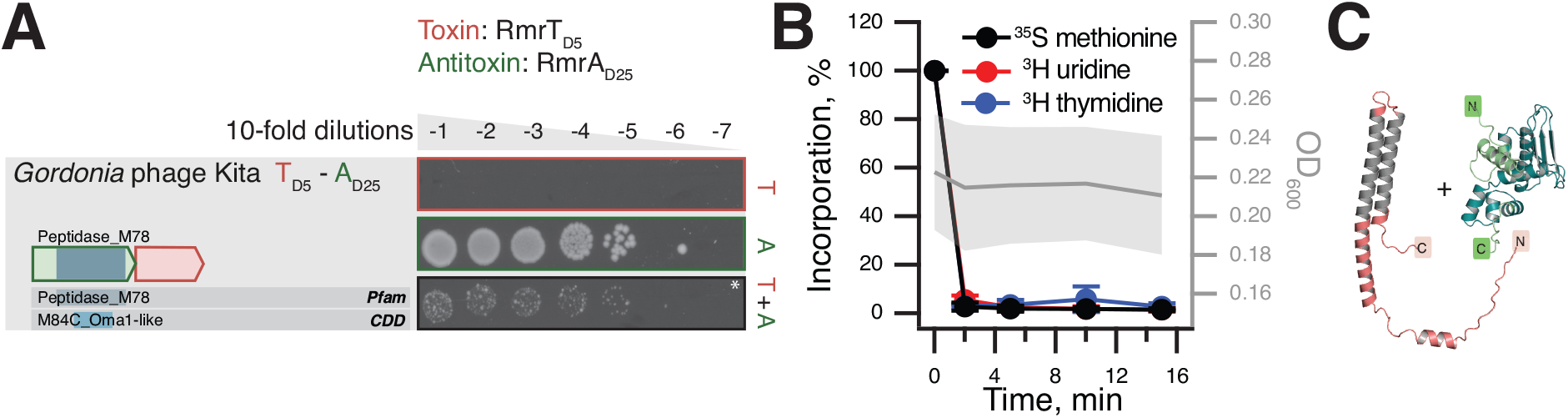
RosmerTA system from *Gordonia* phage Kita. Domain organisation and TA validation though toxicity neutralisation assays (**A**), metabolic labelling assays with toxins expressed in wild-type *E. coli* BW25113 (**B**) and AlphaFold-generated structural models (**C**) for from RmrT_D5_ andRmrA_D25_ TA from Gordonia phage Kita. Predicted transmembrane regions of the toxin are shown in grey. Liquid culture experiments with RmrT_D5_ are shown on **Fig. 4E**.

The TA system from *Acinetobacter guerrae* is composed of a toxin T_D3_ that has no detectable hits with HHPred. However, it has the same fold as mRNA interferases (**Fig. S6**), paired with an AAA ATPase A_D29_ antitoxin (**Fig. 6A**). We refer to the toxin as AarT for <u>A</u>AA-<u>a</u>ssociated <u>R</u>NAse-like <u>t</u>oxin. Metabolic labelling experiments suggest that the toxin targets protein synthesis as its expression inhibits ^35^S methionine incorporation with concurrent increase in ^3^H uridine, a pattern that is characteristic for translation-targeting toxins and antibiotics, and supporting an identity as an mRNAse or tRNase [25]. Similar neutralisation architecture was predicted for D_29_ AAA antitoxins from *Clostridium algidicarnis* (**Fig. 6B**) and *Streptococcus agalactiae* 2603V R (**Fig. 6C**). These two AAA antitoxins are paired with a TOPRIM_OLD domain. The *Lactococcus lactis* AbiL is a bicistronic plasmid-encoded phage system that acts thorough abortive infection elicited by the TOPRIM_OLD toxic effector AbiLii [46]. We speculate that the three AAA-neutralised TA pairs are also Abi phage defence systems. AlphaFold2 modelling does not give a convincing interface for AAA antitoxins and their toxins (pDockQ score of 0.24-0.35), which may be because phage defence AAA-containing systems form large multimeric complexes, as is seen with AAA-containing RADAR [66, 67].

**Figure 6.**
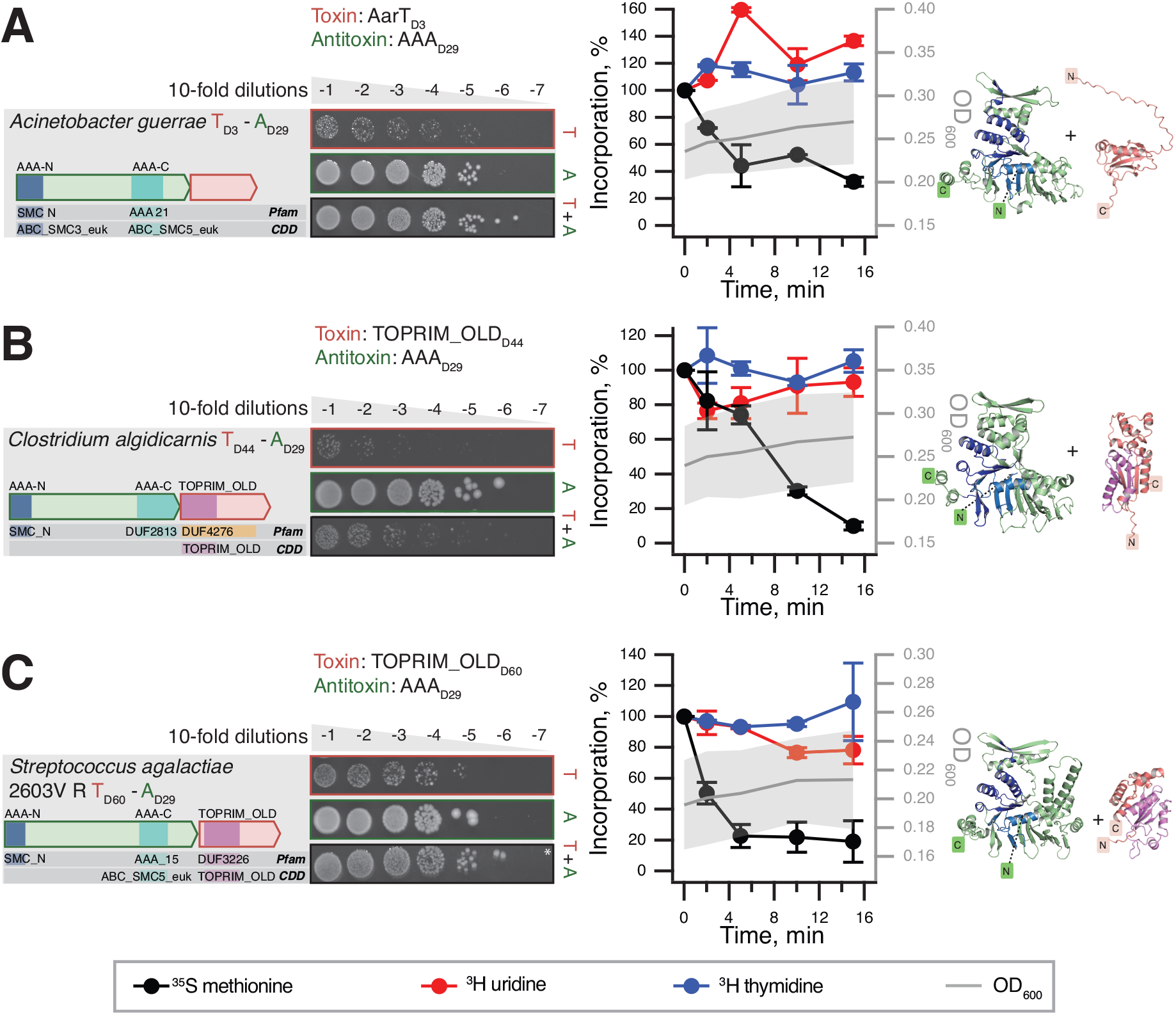
AAA-neutralised putative Abi phage defence systems. Domain organisation and TA validation through toxicity neutralisation assays (*left*), metabolic labelling assays with toxins expressed in wild-type *E. coli* BW25113 (*center*) and AlphaFold-generated structural models (*right*) for AAA_D29_-neutralised TAs: (**A**) *A. guerrae* AarT_D3_:AAA_D29_, (**B**) *C. algidicarnis* TOPRIM_OLD_D44_:AAA_D29_ and (**C**) *S. agalactiae* 2603V R TOPRIM_OLD_D60_:AAA_D29_.

Finally, we have validated 10 TA pairs of nuclease toxins (MqsR_D2_, PIN/VapC-like and YafQ_D119_) paired with diverse antitoxins (PanA, PerlF_D13_, DUF2680_D105_, RHH_6_D107_, RHH_6_D111_, DUF2080_D146_, SynP1_D151_ and RHH_6_D157_), for which AF2 structural models suggest multiple mechanisms of direct and indirect toxin neutralisation (**Fig. 7** and **Fig. S10**,**S11**). While the structures were predicted as binary complexes, RHH (Ribbon-Helix-Helix) domain is a well-characterised dimeric DNA-binding transcriptional regulator employed by numerous antitoxins such as CcdA [68] and FitA [69]. Dimers of RHH-containing antitoxins can be readily predicted by AlphaFold (**Fig. S12**). Multiple groups of translation-targeting RNase TA toxins have been characterised experimentally, and display considerable diversity, even within closely related groups; for example, different VapC PIN TA toxins can cleave either tRNA [70] or rRNA [71]. As expected for nucleases, metabolic labelling assays indicate the majority of the validated nuclease toxins do, indeed, target protein synthesis (**Fig. 7AB** and **Fig. S10**,**S11**). However, unexpectedly, expression of *Thioflavicoccus mobilis* 8321 PIN_D59_ results in inhibition of incorporation of both ^35^S methionine (abrogation of translation) and ^3^H uridine (abrogation of transcription) (**Fig. 7C**). Although metabolic labelling assays with the other two PIN_D59_ toxins (from *Crenothrix polyspora* and *Candidatus Hamiltonella defensa* (formerly *Bemisia tabaci*) do not yield clear-cut results, they are indicative of the PIN_D59_ toxin having additional toxic effects beyond specific inhibition of protein synthesis (**Fig. S10**).

**Figure 7.**
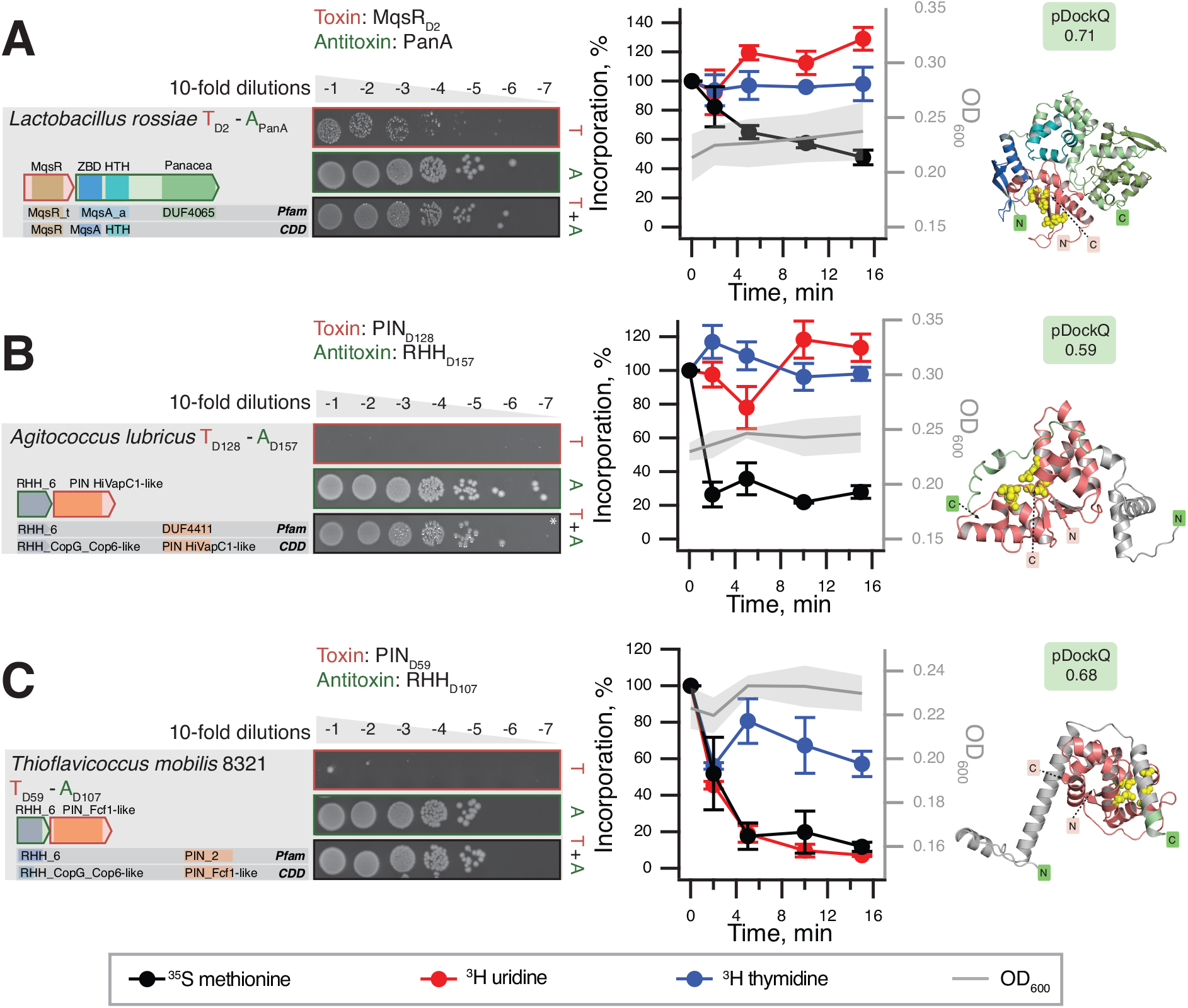
Representative NetFlax TA systems with nuclease effectors: MqsR_D2_, PIN_D128_ and PIN_D59_. Domain organisation and TA validation though toxicity neutralisation assays (*left*), metabolic labelling assays with toxins expressed in wild-type *E. coli* BW25113 (*center*) and AlphaFold-generated structural models (*right*) for TAs with diverse nuclease effectors: (**A**) *L. rossiae* MqsR_D2_:PanA, (**B**) *A. lubricus* PIN_D128_:RHH_D157_ and (**C**) *T. mobilus* 8321 PIM_D59_: RHH_D107_. The active canter of various nucleases in highlighted with yellow spheres on AlphaFold structures.

## Discussion

Classical TA antitoxins are modular proteins typically consisting of a DNA-binding domain involved in transcription autoregulation and a functionally independent neutralisation domain that folds upon binding to the toxin in most cases [1]. It has been argued that it is the combination of, on one hand, the functional decoupling between these two structural modules and, on the other hand, the disordered nature of the neutralisation domains that enables antitoxin promiscuity, i.e. allows for neutralisation of toxins belonging to multiple protein families by different antitoxins possessing the same DNA-binding domain [72]. This model postulates that the fusion of a “linear” recognition motif that performs the neutralisation, along with a DNA-binding domain is sufficient to generate a functional TA operon. Indeed, as we show here for *C. doosanense* Doc_D6_-PanA_Phd-C system, the fusion of the Phd-C region alone to a SUMO tag is sufficient to engineer a protein that efficiently counteract the Phd-C-cognate toxin *in vivo* (**Fig. 3D**). Given the existence of multiple TA operons with single-domain antitoxins that consist of the neutralisation domain alone [73, 74], such fusion or exchange events constitute a plausible evolutionary pathway that could generate the complex toxin-antitoxin permutations observed in TAs (**Fig. 1**).

Importantly, the NetFlax network reveals the existence of a type of hyperpromiscuous antitoxin that defies this commonly accepted neutralisation paradigm. Such antitoxins are epitomised by Panacea and HTH domains [25, 60, 75–77] that possess within their structural fold an intrinsic capacity to specifically recognise and neutralise diverse toxins via three-dimensional epitopes (**Figs. 3C** and **8**). Furthermore, these antitoxin domains can also acquire linear epitopes consisting of intrinsically disordered regions or well-folded domains, to neutralise toxins in the “classical” manner (**Figs. 3EF** and **8**). In addition to this hyperpromiscuity, antitoxins have a capacity for moonlighting in other ways: PAD1-related domains are involved in both toxin neutralisation and phage detection [39], while HigA antitoxins from TAC operons engage both the toxin and the dedicated regulator chaperone [78]. The ability of antitoxins to readily remodel their domain combinations and evolve multifunctional moonlighting abilities of the constituent domains may be a core feature of TA roles in innate immunity involved in phage defence.

**Figure 8.**
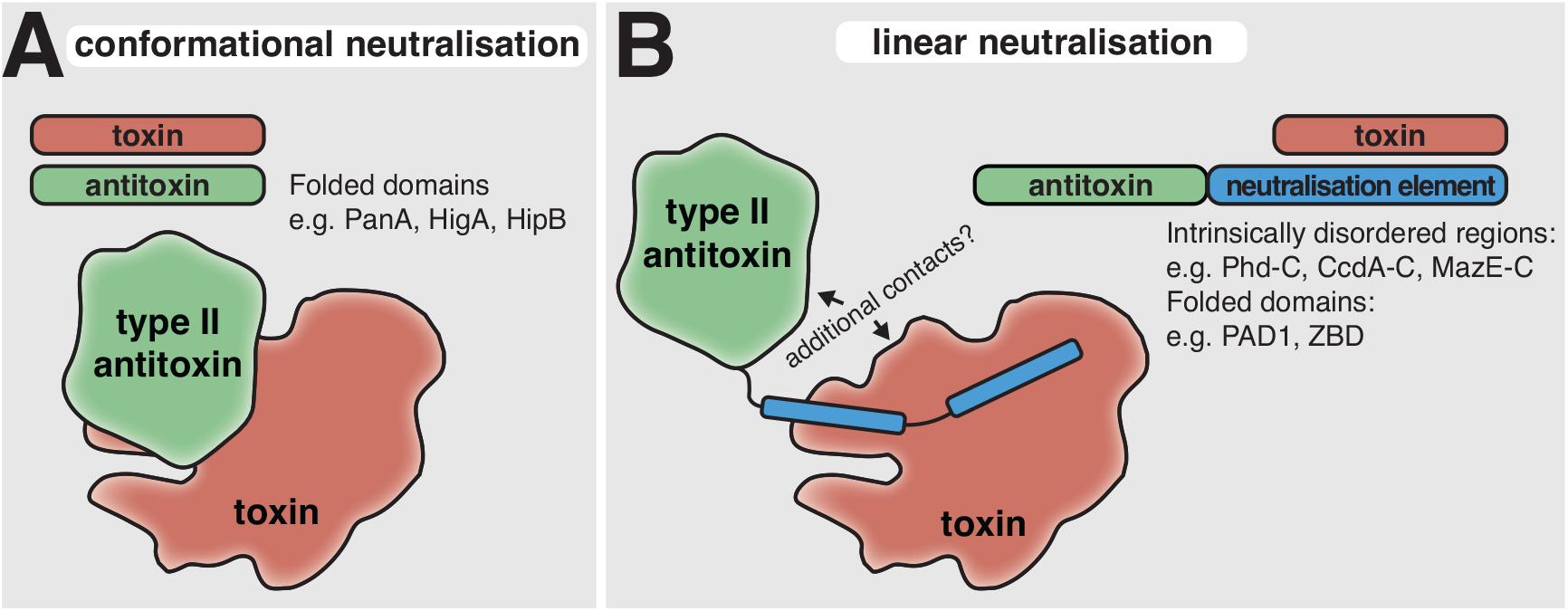
Conformational and linear neutralisation are two primary modes of toxin inactivation in Type II TA systems. (**A**) Conformational neutralisation involves the direct recognition of toxins by a three-dimensional epitope of the antitoxin that forms part of the antitoxin fold and cannot be grafted to a different scaffold. Panacea and HTH domains represent members of this class, with the domains themselves sufficing to neutralise multiple toxins. (**B**) Linear neutralisation displayed by modular antitoxins characterised by the neutralisation of toxins by exchangeable elements that can be intrinsically disordered or fully folded domains. The hallmark of this neutralisation mode is that these neutralising elements are linearly attached to antitoxin domains at the N- or C-terminal ends. The simple permutation of these neutralising elements allows the same antitoxin to neutralise different types of toxins as seen in the case of Panacea linked to Phd_C, ZBD or PAD1, which connects Panacea with the neutralisation of Doc, MqsR and toxSAS toxins. Hyperpromiscuous antitoxins are involved in both types of neutralisation.

In the case of PanAs that function via linear neutralisation without the direct involvement of the globular Panacea domain, the question is what then is the role of the Panacea domain. We hypothesise that its function is primarily sensory, reacting to a trigger and activating the toxin through an allosteric mechanism involving the neutralisation region. This activation may not even require dissociation of the PanTA complex; the fused toxSAS toxin-antitoxin CapRel shows that antitoxins do not have to dissociate in order to activate the toxin [39]. Further investigations are needed to determine what is the functional role of the Panacea domain.

Our structural prediction has allowed the clustering of predicted toxins and antitoxins into identifiable fold classes. This tendency of TA systems to reuse folds has previously been noted [60]. However, conservation of fold may not mean conservation of function: the proteins can be divergent at the sequence level – even to the point of carrying out different biochemistry, despite being based on the same structural fold. For example, the Fic/Doc family of toxins contains members that can both NMPylate or phosphorylate their protein targets [63, 79]. Similarly, toxSAS TAs can either inhibit bacterial growth by producing toxic alarmone (pp)pApp or pyrophosphorylating the 3’ CCA end of tRNA [19, 80]. Our *T. mobilis* PIN_D59_ domain toxin that seems to inhibit transcription as well as translation may indicate another example of a divergent function on a similar fold.

The biological function of TAs has remained a contentious subject for decades [1, 2, 81], but increasingly a role for these systems in phage defence is being discovered [12, 14, 35–38]. Here we show that this is also reflected in their evolutionary relationships: the core network of TAs is connected to domains other phage defence systems such as PARIS, AbiLi, Septu, Gabija and Rosmer systems [13, 35, 46, 47, 50]. Indeed, since it is common for multigene defence systems to contain a toxic effector, it is unclear whether there is any meaningful distinction between the two kinds of system. This raises the question of whether classical “addiction module” TAs on mobile elements may actually have a role in defence against phages or competition with other mobile elements, with addiction effects being a secondary consequence.

NetFlax is a broad stroke approach, which has its limitations and caveats that we acknowledge. First, it is limited to a representative set of proteomes, which means we are missing a substantial amount of diversity. Some known Type II TAs such as DarTG [82], HEPN-MNT [83], HicBA [84], and HipBA [85] escaped our prediction. Second, NetFlax only addresses conserved two-gene proteinaceous systems and therefore can not in this incarnation predict multi-gene toxin containing systems. It is also at risk of predicting false positives due to spurious domain associations. Nevertheless, despite these caveats, the network is a starting point for exploring multiple new avenues including the 20 TA systems we have characterised, and more fine-grained prediction can be achieved through a subsequent focus on specific lineages.

## Materials and methods

### NetFlax strategy

TA pairs were predicted with the Python script NetFlax, which is a modification of our FlaGs program. The overall strategy is based on the pipeline of **Fig. S1**. Briefly, NetFlax works round by round; and each round follows three basic steps: i) scanning of a local database of proteomes using an HMM profile of toxin or antitoxin to identify its protein homologues, ii) prediction of conserved TA-like arrangements by analysing the neighbourhood of the protein homologues and identification of homologous clusters of toxins or antitoxins and iii) cross-checking if the predicted clusters are novel (never been identified in any previous round) and, if so, make their HMM profiles, to be used in the next round of scanning mentioned in step 1.

NetFlax uses a local database of 24,479 predicted proteomes downloaded from the NCBI RefSeq FTP server [86]. The local database includes one representative proteome per species of bacteria and archaea, along with all 10,449 available virus genomes (not limited to representatives). NetFlax uses conservation of gene neighbourhood analyses by FlaGs and combines it with the guilt-by-association strategy to identify TA-like pairs. NetFlax is multi-core aware and proceeds round by round hopping from antitoxin to toxin and toxin to antitoxin and continuing similarly until it finds the entire network (see **SI Text, Fig. 1** and – in more detail – **Fig. S1** for illustrations of how the algorithm works).

### Protein function prediction

Protein domains and other functional predictions were carried out with searching the toxin-antitoxin database (TADB) [48], DefenceFinder [43], NCBI conserved domain database (CDD) [87] and HHPred [88] (with NCBI-CDD, Pfam-A and PDB as target databases).

### Protein structure prediction and comparisons

Protein-protein complex structures were predicted with FoldDock [45]. Structures were clustered with FoldSeek v. 1.3 [89] and with resulting networks visualised with Cytoscape v. 3.5.0 [90]. Structural alignment was carried out with mTM-Align v. 20220104 [91]. Transmembrane prediction was carried out with DeepTMHMM [92]. More detailed methods are described in the **SI Text** document.

### Experimental methods

Detailed experimental procedures are provided the **SI Text** document, with a summary below.

#### Plasmid construction

All bacterial strains, plasmids and primers used in the study are listed in **Dataset S2**. Toxin ORF-s were cloned into arabinose-inducible pBAD33 vector [93] either with or without Shine-Dalgarno sequence as required, for toxicity assay. For neutralization assay, antitoxins were expressed from an IPTG inducible pMG25 vector [94]. Mutations and truncations were introduced to the antitoxin genes as described earlier [23].

#### Toxicity neutralisation assays

Toxicity-neutralization assays were performed on Lysogeny broth (LB) agar plates. First, the pBAD33 vector with the toxin ORF was transformed into competent cells of *E. coli* BW25113 strain with a pMG25 empty vector. A single colony with two plasmids was grown in liquid LB medium supplemented with 100 μg/mL ampicillin (Sigma-Aldrich) and 20 μg/mL chloramphenicol (AppliChem) as well as 0.2% glucose (repression conditions). Serial 10-fold dilutions were spotted (5 μL per spot) onto solid LB plates containing ampicillin and chloramphenicol under repressive (0.2% glucose) or induction conditions (0.2% arabinose combined with 0.05 or 0.5 mM IPTG). Plates were scored after an overnight incubation at 37 °C. After confirming toxin toxicity, another set of competent cells was produced with the cognate antitoxin in a pMG25 vector, and the spot test was repeated with all possible combinations.

#### Metabolic labelling

metabolic labelling experiments using *E. coli* BW25113 strains co-transformed with pBAD33 derivatives as well as the empty pMG25 vector were performed as described earlier [25].

## Acknowledgments

We are grateful to the Protein Expertise Platform at Umeå University for plasmid construction. The AlphaFold2/FoldDock computations were enabled by the supercomputing resource Berzelius provided by the National Supercomputer Centre at Linköping University and the Knut and Alice Wallenberg foundation. This work was supported by Knut and Alice Wallenberg Foundation (2020.0037 to GCA), the Swedish Research council (2019-01085 and 2022-01603 to GCA, 2021-01146 to VH, 2021-03979 to AE); Crafoord foundation (project grant Nr 20220562 to VH); Cancerfonden (20 0872 Pj to VH); Carl Tryggers Stiftelse för Vetenskaplig Forskning (CTS19:24 to GCA); the European Regional Development Fund through the Centre of Excellence for Molecular Cell Technology (VH and TT); Ragnar Söderberg foundation (VH); Fonds National de Recherche Scientifique (FNRS CDR J.0068.19, FNRS-EQP UN.025.19 and FNRS-PDR T.0066.18 to AG-P); ERC (CoG DiStRes, n° 864311 to AG-P).

