## Supplementary material for "The structural basis of hyperpromiscuity in a core combinatorial network of Type II toxin-antitoxin and related phage defence systems": SI text and figures

### Supplementary methods

*NetFlax sequence searching*
For sequence searching we used a local proteome database of 24,479 predicted proteomes downloaded from the NCBI Genomes FTP server (ftp.ncbi.nlm.nih.gov/genomes) on the 29th of January, 2021, selecting all viruses, and one representative proteome per species for archaea and bacteria (1, 2).

In each round, NetFlax follows three steps, which are summarised here and described in detail in the sections below: (1) find protein homologues of toxins and antitoxins across the input genomes using Hidden Markov Model (HMM) profile-based homology sequence similarity searching (2) assessing gene neighbourhood conservation of the identified homologues to predict TA-like arrangements and novel clusters of potential toxin and antitoxin candidates (3) "hop", that is repeat step 1, using the HMM profiles of novel clusters identified (**Fig. 1**). The final step (4), is the summation of results as a network. All scripts and datasets are available at https://github.com/GCA-VH-lab/coreNet.

*Step 1: HMM searching*
NetFlax uses HMMsearch package from HMMer v3.1b2 for scanning the local proteome database to identify homologues to complete *Step 1* (3). To start, we used the HMM profile of DUF4065/Panacea domain downloaded from the Pfam database (4), that has been recently recognized as hyperpromiscuous antitoxin domain (5). The e-value threshold for homology searching is generally set as $1e^{-10}$, with the exception for DUF4065/Panacea, which uses the Pfam-set HMM profile's score-based curated gathering threshold (4). At the end of each homology searching, all the identified protein homologues including their corresponding HMM profile were stored in a database for use in the subsequent cross-checking steps of the NetFlax pipeline (see **Fig. S1**).

*Step 2: predicting TA-like architectures*
For *Step 2*, the identified protein homologues from each profile are filtered to create input files for the analysis of gene neighbourhoods using an adapted version of FlaGs (see **Fig. S1**) (6). The filtering criteria for making the input file were as follows: (1) the length of the protein should not be more than 450 amino acids, and (2) the input file should not carry more than 1,000 protein accessions, at which point the procedure becomes too computationally challenging. When the number of identified protein homologues exceeds the limit of 1,000 accessions, the NetFlax pipeline uses Usearch v11.0.667 to create a reduced representative set of sequences from the initial number of identified protein homologues (see **Fig. S1**) (7). We applied a cut-off value that ranges from 0.1 to 0.9 for the '-id' argument while performing analyses with Usearch. With the Usearch option -id 0.9, a list of sequences is generated that are representatives of 90% identical clusters, while with option -id 0.1 it produces a list of sequences that are representatives of 10% identical clusters. Using a range of values from 0.1 to 0.9 we were able to select representative sequences that are less or equal to 1000. While this means that some pairs did not always make it through to the next round, they were retained in the list of final hits reported, and representative relatives went through to the next round.

The accessions from the identified – and if necessary filtered - sequences are used to make the input file for the prediction of TA-like arrangements (**Fig. S1**). The code for this comes from FlaGs, which takes advantage of the sensitive sequence search method Jackhmmer (3), to cluster neighbourhood-encoded proteins into homologous groups (6). NetFlax filters TA-like arrangements based on the following criteria: (1) the genes in the predicted TA-like arrangement should be in the same order; (2) the intergenic distance between two genes should not exceed 100 nucleotides; (3) the conservation of the gene neighbourhood should be limited to the two genes that are in close proximity (thus excluding longer conserved operons); (4) the length of the predicted toxin or antitoxin should not be more than 450 amino acids; and (5) the identified TA-like architecture should be conserved in a certain number of species/taxa across the dataset; which we defined as the taxa limit. We tested 6, 8 and 10 as taxa limits and compared the results (see **Dataset S1**). Gene pairs that met those criteria were subjected to another filtering step that we call cross-checking (see **Fig. S1B**).

NetFlax applies a sequence similarity searching approach that involved scanning of each identified potential cluster against the cross-checking database to determine whether the cluster is novel. This database contains all the protein accessions identified from the HMM-profile based homology searching steps mentioned above. Also, it includes all the HMM profiles, and the database is continuously updated throughout the NetFlax rounds until hopping ends. Inter-round hit in the cross-checking step ends up as back crosses (dotted lines in **Fig. 1**) in the resulting network. The filtering process of finding novel clusters involved two steps (**Fig. S1B**). First, the protein accessions in the identified clusters were checked to determine if they were found in the cross-check database. If a similar accession was discovered, the detected cluster was not considered novel and hence did not advance to the next round. Otherwise, if the detected cluster passed the initial phase of cross-checking, the protein sequences from the cluster were submitted to HMM-profile based similarity searching (as a second step of filtering), which used the HMM profiles stored in the cross-check database. The e-value threshold was set as in *Step 1*. If any of the previously stored HMM profiles found a hit while scanning against the protein sequences from a cluster, the cluster did not proceed to the next round of hopping.

We found that lower taxa limits led to large networks that upon inspection contained many lineages in the outer parts of the network comprised of relatively large gene pairs that were rarely overlapping and were indicative that NetFlax had wandered beyond the realms of toxin-antitoxins and related systems. For example, when we decreased this taxa limit to 6, we identified 26,644 TA-like pairs that are clustered into 1,556 homologous groups of putative toxins and antitoxins, well beyond our ability to validate (see **Table Sx**). While some false positives are to be expected, we chose on balance to be more restrictive to limit their overrepresentation and compute an initial core network that we are more confident in, and where it is possible to experimentally validate most novel domain pairs and predict all structures. A downside of this stringency is that some TAs inevitably go under the radar. Therefore, we added a final guilt by association hop that only required a taxa limit of two (*See Step 4*, below).

*Step 3: Hopping to the next round*

When a potential cluster passes both steps of cross-checking (see **Fig. S1B**), protein sequences involved in that cluster are then retrieved and aligned using MAFFT v7.453 with the L-INS-i strategy (8). Then, a new HMM profile was made from that alignment using the hmmbuild package of HMMer v3.1b2 (see **Fig.S1**) (3). The new HMM profiles built from novel identified clusters can then to hop to the next round and the cross-checking database is updated (see **Fig. S1**). It is possible that NetFlax may find multiple new HMM profiles from a single round (see **Fig. S1**). Therefore, NetFlax processes multiple tasks in parallel.

*Step 4. Reducing the stringency of criteria to predict additional nodes*

In order to find additional homologues at this point, we carry out a search that only requires the two-gene architecture to be conserved in two genomes. These results are reported, but if the conservation across fewer than eight genomes, the hits do not proceed to a subsequent hop, for concern of the greater risk of false positives when reducing the conservation stringency.

*Step 5: Visualisation of TA-like clusters as a network*

We used a python script that uses a module called pyvis (https://pyvis.readthedocs.io/en/latest/) to make an interactive network of all the predicted toxin and antitoxin clusters annotated based on the predicted domain they are carrying (discussed below). The interactive network is available at: http://netflax.webflags.se/. The clusters of predicted toxins are represented as red circles, whereas predicted antitoxins are shown in blue circles. Cytoscape was used to create a vector graphic image of the TA network across all micro-organisms (see **Fig. 2**) (9).

*Annotation of the predicted toxin and antitoxin clusters*

To annotate the predicted toxin and antitoxin clusters in the interactive network, one representative protein sequence from each cluster was checked if they share identifiable homology with any known domain in the National Center for Biotechnology Information Conserved Domains Database (NCBI CDD) (10); and to any known structure or protein family with any probability greater than 50%, that can be searched with HHpred by selecting the databases of protein families (Pfam) and protein data bank (PDB) (11). Furthermore, Blastp was used to assess the sequence similarity of selected representative sequences from each predicted cluster to the experimentally confirmed type II toxin and antitoxin protein sequences from TADB 2.0 (12). In order to do that, we built a reference sequence database using the downloaded protein sequences from TADB of toxins and antitoxins. To determine similarity with Blastp, the e-value cut-off was set to 1e$^{-3}$, with an additional parameter '-qcov_hsp_perc', which was set to 45. The '-qcov_hsp_perc' parameter allows to prevent spurious alignments of a short segment of the query sequence to the reference sequences stored in the database. At least 45% of the representative sequence from each predicted cluster has to form an alignment with the toxin or antitoxin sequences in the reference sequence database with an e-value threshold of 1e$^{-3}$ to be considered a hit. Similarly, we conducted Blastp analyses in which the reference sequence database was made up of sequences from previously verified toxSAS-antitoxSAS TA pairings (13). The search for domains associated with phage defence systems was performed using hmmscan with an E-value 1e$^{-3}$ cut-off features from HMMER package (version 3.3.2) with the DefenseFinder HMM profile database (downloaded in May 2022).

We added an additional HHPred-based annotation step for the summary information in **Dataset S1**. We predicted the domains of all proteins in each node in order to uncover the most common domain and PDB hits. These are reported as percentages for each node.

*Identification of the core proteinaceous TA network*

NetFlax applies several filtering criteria to compute the core network of proteinaceous TA systems. One limit is that none of the proteins should be longer than 450 amino acids. To date, the longest toxin reported in the TADB database of the HipB/HipA system is 440 amino acids, and antitoxins are usually smaller (12). Therefore, we decided that the length of the protein sequence predicted as toxin or antitoxin should not exceed a limit of 450 aa. Other criteria used in the prediction of TA like arrangements are that the two adjacent genes should be conserved in the same order in a region that is otherwise not conserved, and should not have an intergenic distance of more than 100 nucleotides. Also, the predicted TA-like architecture should be found conserved at least in a certain number of species/taxa across the dataset. We call this the taxa limit. For example, taxa limit 10 means that each of the predicted TAs should be conserved in at least 10 taxa across the dataset. With lower limits, the chance that we are discovering false positives also increases and is at a scale where experimental validation of lineages becomes unfeasible. We chose a taxa limit of 8 results, which produced a non-exhaustive but large network, still small enough to examine by eye, predict structures, and validate lineages experimentally.

*Protein complex prediction*

The simultaneous folding and docking protocol FoldDock (14) was implemented to predict the heterodimeric Toxin-Antitoxin (TA) complexes from the NetFlax network. The FoldDock protocol predicts dimers using optimized multiple sequence alignments and passing the paired alignments through the AlphaFold2 pipeline. Based on the number of residues and quality of the interface, a pDockQ (predicted DockQ) score is calculated (Python script available from https://gitlab.com/ElofssonLab/huintaf2/-/blob/main/bin/analysepairs.py, 10Å used as the interface distance cut-off). High scoring acceptable models are usually considered at DockQ ≥ 0.23 (14). For selecting complexes for experimental validation of interfaces, we have been more conservative, only considering pDockQ scores of ≥ 0.4 as strongly indicating interacting surfaces. The FoldDock protocol was run to score multiple models (5 models per pair) and the predicted complex with the highest pDockQ score was selected to represent the TA pair complex structure. A single representative AF2 model from each node in the NetFlax network was selected to build a representative dataset of 320 structures (consisting of both toxins and anti-toxins as per the node distribution) in order to perform clustering analysis. We constructed the representative dataset by selecting an AF2 model based on either a predicted complex which has been experimentally validated by us in the lab or by considering a predicted complex with the highest available PdockQ score (14) in case the experimental data is not available. This filtering process was repeated for all the nodes present in the NetFlax mininode network to build the representative dataset of the 320 predicted AF2 models.

*Structural clustering*

A single representative model from each node in the NetFlax network was selected to build a representative dataset of 320 predicted structures (consisting of both toxins and anti-toxins according to the node distribution) in order to perform clustering analysis. The predicted model for each node was selected either based on experimental validation confirmed by us in the lab or by considering a predicted complex with the highest available PdockQ score (14) in case the experimental data is not available. This filtering process was repeated for all the nodes present in the NetFlax network to build the representative dataset (as listed in **Dataset S1**).

Foldseek (v5, (15)) was run to conduct a protein structure similarity search within our structural dataset to find clusters of proteins with similar folds. The searches were conducted based on the Gotoh-Smith-Waterman local alignment by implementing the easy-search as well as the cluster modules available and the resulting output was utilized to construct a structural similarity network (SSN) with the help of Cytoscape program (v3.5.0, (16)) based on the E-value. In the Cytoscape program the SSN was visualized by implementing the organic layout option with a node filter based on E-values below $1e^{-3}$.

Subsequently, we used Foldseek (15) to conduct a structural homology search within the aforementioned AF2 dataset to elucidate any underlying clustering patterns and visualize it in the form of a structural similarity network (SSN). Herein, the Foldseek program searches were performed based on the Gotoh-Smith-Waterman local alignment and the resulting output i.e. the E-values were utilized to construct the SSN and visualize the clustering patterns (between the AF2 model dataset) with the help of Cytoscape program (16). Next, the resulting network for our representative dataset was visualized by implementing the organic layout option (available in the yFiles Layouts option) with a node filter based on E-values below $1e^{-3}$. This results in the formation of total seventeen clusters with eight larger clusters (with three or more constituent nodes) along with nine smaller clusters (less than three constituent nodes). We also performed structural alignment of the major clusters using the mTM-alignment program (17). Finally, we mapped these clusters on to the Netflax network.

*Structural alignment of toxSAS toxins*
To align PAD1 domain in neutralizing toxSAS toxins, we used the DALI Protein Structure Comparison Server with pairwise alignment (18). Our query was the PAD1 domain from *Clostridium hylemonae* DSM 15053 $PAD1_{D27}$, which was aligned with *B. subtilis* Ia1a PanA (only the first 94 aa used) and fused TA CapRel from *Salmonella* phage SJ46 (272-373 amino acids) (19). The DALI run indicated a similar fold with a Z-score > 3.5.

*Transmembrane topology prediction*
Transmembrane regions were predicted with DeepTMHMM (20).

*Construction of plasmids*
To test toxicity, toxin ORFs were cloned into the medium copy, arabinose inducible pBAD33 vector (21) using Gibson assembly (22) or the CPEC protocol (23). In Gibson assembly the vector was linearised with primers VTK26 and VTK58 (adding a strong Shine-Dalgarno (SD) motif and the intervening sequence 5'-AGGAGGAATTAA-3'). When expression of the toxin caused a growth defect to the host *E. coli* BW25113 in repressed conditions, the SD region was deleted using VJD15 and VJD16 primers. The toxin containing ORF was chemically synthesized by IDT with flanking regions of the plasmid (5'-TCGAGCTCAGGAGGAATTAA-3' or 5'-CCGCCAAAACAGCCAAGCTT-3'), SD region underlined. Cloning of constructs without SD is shown in **Dataset S2** using the version with Shine-Dalgarno as a template.

To test the neutralisation of the toxin a high copy, IPTG inducible, pMG25 vector was used (24) for antitoxin expression. All the constructs were constructed with a strong SD sequence. For Gibson assembly the plasmid was linearized with PCR primers VJD1 and VJD2. Synthetic DNA harbouring the ORF of the antitoxin was ordered with flanking regions of the plasmid backbone (5'-CCAAGCTTAGGAGGAATTAA-3' and 5'-TCAACAGGAGTCCAAGCTCA-3') and cloned to pMG25 vector with Gibson assembly as shown in **Dataset S2**. In some cases, an antitoxin was sub-cloned from the pKK223-3 (25) plasmid to pMG25 vector using long primers suitable for Gibson assembly (**Dataset S2**).

Plasmids VHp1101, VHp1102, VHp1124 and VHp1125 were constructed following Circular Polymerase Extension Cloning (CPEC) protocol (23). The toxin containing ORF was cloned to pBAD33 vector resulting with plasmids VHp1101 and VHp1124. For that, pBAD33 vector was linearized with restrictases *Sac*I and *Hind*III and synthetic DNA with flanking regions (5'-CGAATTCGAGCTCAGGAG-3' and 5'-CTCATCCGCCAAAACAGCCAAG-3') was ordered from IDT. The ORF containing fragment was amplified with PCR primers pBAD_SD_TOX_fwd and pBAD_TOX_MCS_rev. Plasmids VHp1102 harbouring an antitoxin was constructed first with pKK223-3 backbone: medium copy number, ColE1 origin of replication, $Amp^R$, antitoxins are expressed under the control of a $P_{Tac}$ promoter (25). For that *Eco*RI and *Hind*III were used to linearize the plasmid and ORF of the antitoxin was ordered from IDT with flanking regions (5'-CGAATTCGAGCTCAGGAG-3' and 5'-CTCATCCGCCAAAACAGCCAAG-3'). The ORF containing fragment was amplified with PCR primers pKK_ATOX_MCS_fwd_2 and pKK_ATOX_MCS_rev2 and inserted to pKK223-3 backbone with CPEC. To obtain the same level of antitoxin expression, only pMG25 vector was used in neutralisation assays. For that long primers compatible for Gibson assembly or CPEC were ordered and antitoxin from VHp1102 was cloned to a new vector resulting with VHp1371. Plasmids VHp1125 was constructed with pMG25 backbone harbouring an antitoxin ORF with CPEC (see **Dataset S2**).

Mutations and truncations to the gene of the antitoxin were introduced as described earlier (26). Single-point mutations were introduced to PanA antitoxins with Phusion DNA Polymerase (Thermo Scientific) by amplifying the whole plasmid with one of the primers was carrying the desired mutation. For deletion mutations, primers were designed to insert an early stop-codon or to introduce start-codon if necessary (**Dataset S2**). After PCR amplification, the product was treated with *Dpn*I, purified with Monarch PCR & DNA Cleanup Kit (NEB), treated with T4 Polynucleotide Kinase (Thermo Scientific) and ligated with T4 DNA Ligase (Thermo Scientific). The mixture was transformed to *E. coli* DH5α strain. Plasmids were purified with E.Z.N.A. Plasmid DNA Mini Kit (Omega) and verified by sequencing.

*Metabolic labelling*

For metabolic labelling experiments, *E. coli* BW25113 strains co-transformed with pBAD33 derivatives (for L-arabinose-inducible expression of toxins) as well as the empty pMG25 vector were first plated out on LB plates supplemented with 100 µg/ml carbenicillin, 25 µg/ml chloramphenicol and 0.2% glucose (to suppress the leaky expression of the toxin). Using fresh, individual *E. coli* colonies for inoculation, 2 mL liquid cultures were prepared in defined Neidhardt MOPS minimal media (27) supplemented with 100 µg/ml carbenicillin, 25 µg/ml chloramphenicol, 0.1% of casamino acids, and 0.2% glucose, and grown overnight at 37 °C with shaking. Next, experimental 15-mL cultures were prepared in 125 mL conical flasks in MOPS medium supplemented with 0.5% glycerol, 100 µg/ml carbenicillin, 25 µg/ml chloramphenicol as well as a set of 19 amino acids (lacking methionine), each at final concentration of 25 µg/mL. These cultures were inoculated overnight to final $OD_{600}$ of 0.05 and grown at 37 °C with shaking up to of $OD_{600}$ 0.2. At this point, one 1-mL aliquot (the pre-induction zero time-point) was transferred to 1.5 mL Eppendorf tubes containing 10 µL of radioisotope – either $^{35}S$ methionine (4.35 µCi, Perkin Elmer), or $^{3}H$ uridine (0.65 µCi, Perkin Elmer) or $^{3}H$ thymidine (2 µCi, Perkin Elmer) – and transferred to the heat block at 37 °C. Immediately after, the expression of toxins in the remaining 14 mL culture was induced by addition of L-arabinose (final concentration of 0.2%). Throughout the toxin induction time course, 1-mL aliquots were taken from the 15 mL culture and transferred to 1.5 mL Eppendorf tubes containing 10 µl of radioisotope ($^{35}S$ methionine / $^{3}H$ uridine / $^{3}H$ thymidine). The incorporation of radioisotopes was stopped after 8 minutes of incubation at 37 °C by adding 200 µL of ice-cold 50% trichloroacetic acid (TCA) to 1 mL cultures. In parallel with taking the time-points for labelling, 1 mL aliquots were taken for $OD_{600}$ measurements. Isotope incorporation was quantified by normalising radioactivity counts (CPM) to $OD_{600}$, with the pre-induction zero time-point set as 100%. All experiments were performed in three biological replicates (i.e. using three independent cultures inoculated from three different colonies).

*Growth assays*

Growth assays were performed in liquid MOPS medium (1x MOPS mixture (AppliChem), 1.32 mM $K_2HPO_4$ (VWR Lifesciences), 1 mg/ml thiamine (Sigma), 25 µg/ml each of 20 amino acids and the carbon source – either 0.5% glycerol (VWR Chemicals) or 1% glucose. The media was supplemented with ampicillin and chloramphenicol. Overnight cultures were grown in MOPS medium supplemented with 1% glucose at 37 °C. The cultures were diluted to a final $OD_{600}$ of 0.05 in MOPS medium supplemented with 0.5% glycerol and then growth was monitored using a Bioscreen C Analyzer (Oy Growth Curves Ab Ltd) at 37 °C for 10 hours. Toxins were induced at around $OD_{600}$ = 0.4 with 0.2% arabinose.

| Organism | Antitoxin | | | Toxin | | |
|---|---|---|---|---|---|---|
| | Name | Accession | Domains present | Accession | Domains present | Name |
| *Acinetobacter guerrae* | $AAA_{D29}$ | WP_152878174.1 | AAA-N; AAA-C | WP_152878172.1 | None predicted; RNA-like fold | $AarT_{D3}$ |
| *Agitococcus lubricus* | $RHH_{D157}$ | WP_107864200.1 | RHH_6 | WP_107864201.1 | PIN_HiVapC-like | $PIN_{D128}$ |
| *Bacillus fungorum* | PanA | WP_099686064.1 | Panacea; ZBD (MqsA); HTH | WP_099686063.1 | MqsR | $MqsR_{D2}$ |
| *Bacillus subtilis* la1a | PanA | WP_090558408.1 | Panacea | WP_090558406.1 | RelA/SpoT (ToxSAS) | $PhRel2_{D8}$ |
| *Bartonella choladocola (Bartonella apis)* | PanA | WP_077993240.1 | Panacea | WP_077993242.1 | Transmembrane | $PanT_{D11}$ |
| *Candidatus Hamiltonella defensa (Bemisia tabaci)* | $RHH_{D111}$ | WP_016857969.1 | RHH_6 | WP_016857968.1 | PIN_Fcf1-like | $PIN_{D59}$ |
| *Clostridium algidicarnis* | $AAA_{D29}$ | WP_185162150.1 | AAA-N; AAA-C | WP_104409920.1 | TOPRIM_OLD | $TOPRIM\_OLD_{D44}$ |
| *Clostridium hylemonae* DSM 15053 | $PAD1_{D27}$ | WP_006441954.1 | PAD1 | WP_138261968.1 | RelA/SpoT (ToxSAS) | $PhRel2_{D8}$ |
| *Corynebacterium doosanense* | PanA | WP_018022803.1 | Panacea | WP_018022804.1 | Fic/Doc | $Doc_{D6}$ |
| *Crenothrix polyspora* | $SpnP3_{D105}$ | WP_087148009.1 | DUF2680 | WP_087148008.1 | PIN_VapC-like | $PIN_{D59}$ |
| *Desulfonatronovibrio magnus* | $SynP1_{D151}$ | WP_045215642.1 | Ribosomal L12_N; unstable_antitoxin; DinJ-like | WP_045215644.1 | YafQ toxin | $YafQ_{D119}$ |
| *Gelidibacter mesophilus* | PanA | WP_027127283.1 | Panacea | WP_027127282.1 | FtsL; transmembrane | $toxFtsL_{D9}$ |
| *Gordonia* phage Kita | $RmrA_{D25}$ | YP_009301393.1 | Peptidase_M78 (ImmA/RmrA) | YP_009301394.1 | Transmembrane | $RmrT_{D5}$ |
| *Lactobacillus rossiae* | PanA | WP_017263061.1 | Panacea; ZBD (MqsA); HTH | WP_017263062.1 | MqsR | $MqsR_{D2}$ |
| *Mucilaginibacter gotjawali* | $SpnP1_{D153}$ | WP_157750404.1 | None known | WP_096349837.1 | DUF4411/PIN_VapC | $PIN_{D128}$ |
| *Streptococcus agalactiae* 2603V R | $AAA_{D29}$ | WP_000491711.1 | AAA-N; AAA-C | WP_000010056.1 | TOPRIM_OLD | $TOPRIM\_OLD_{D60}$ |
| *Thioflavicoccus mobilis* 8321 | $RHH_{D107}$ | WP_015281565.1 | RHH_6 | WP_015281564.1 | PIN 2 | $PIN_{D59}$ |
| *Thiomicrospira cyclica* ALM1 | $SpnP2_{D146}$ | WP_013835461.1 | DUF2080 | WP_013835460.1 | DUF4411/PIN | $PIN_{D128}$ |
| *Thiorhodococcus minor* | $PrlF_{D13}$ | WP_164450688.1 | PrlF | WP_164450689.1 | DUF4411/PIN | $PIN_{D54}$ |

**Table S1. NetFlax TAs that were experimentally characterised in this study.**

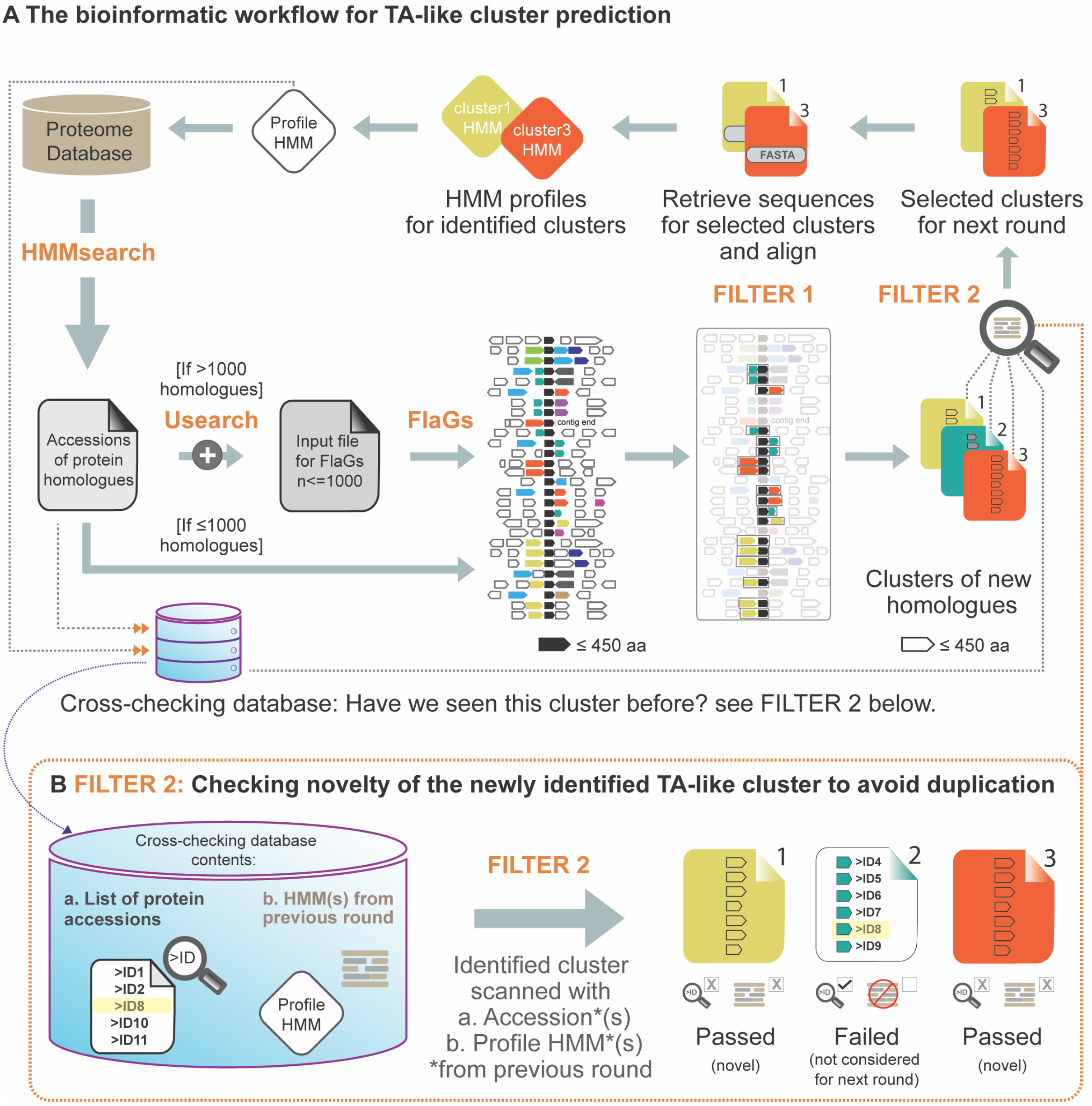


**Figure S1. The bioinformatics workflow for TA-like cluster prediction.**
(**A**) NetFlax process begins with scanning our local proteome database using an HMM profile (for a start, we used an HMM of Panacea domain) to identify protein homologues. All the protein homologues identified from the scanning, including the HMM profile, constitute the cross-checking database used later to identify novel clusters of potential toxin and antitoxin candidates. Selected 1000 protein homologues (each with a length no longer than 450 amino acids) are then used for neighborhood conservation analyses using an adapted version of FlaGs. FlaGs automatically predicts the two-gene architectures that are typical of TA loci using the following filtering criteria:
1. Two components of the predicted TA arrangement should be in the same order and does not have an intergenic distance of more than 100 nucleotides
2. The maximum length of the predicted toxin or antitoxin should not exceed 450 amino acids; and 3. The predicted TA-like architecture should be found at least conserved in eight species/taxa across the dataset.
The identified clusters of partner genes from FlaGs analyses were then filtered based on a sequence similarity searching approach (details in C) to ensure that the identified clusters were distinct. When the novel potential clusters of toxin or antitoxin candidates are found, NetFlax makes HMM profiles specific for each cluster. These HMM profiles were then used for the next round of hopping.
(**B**) Further detail of the FILTER 2: Checking novelty of the newly identified TA-like cluster to avoid duplication. FILTER 2 testing depends on the cross-checking database, which is continuously updated throughout the NetFlax process with the identified protein homologues and their corresponding HMMs

from each hopping round. When the potential cluster is found from the FlaGs analyses, it is then subjected to the FILTER 2 test includes two following steps:

1. 1st step involves checking if any accession in the potential cluster is already present in the cross-checking database. If found, that means the cluster is homologous to a previously found cluster. Therefore, to avoid duplication of the homologous cluster in the network we do not allow this cluster to proceed to further steps of the NetFlax pipeline. If not found, we consider the cluster for the next filtering step described below.
2. If the potential cluster passes the 1st step, we scan all the protein sequence of the cluster using all the HMM profiles present in the cross-checking database to detect homology. If homology is detected with any of the HMM profiles present in the cross-checking database, the cluster is then considered as failed and therefore not allowed for the next pipeline steps.

If the potential partner gene cluster passes both steps in FILTER 2, it is considered novel and used to make a new HMM profile for the next round.

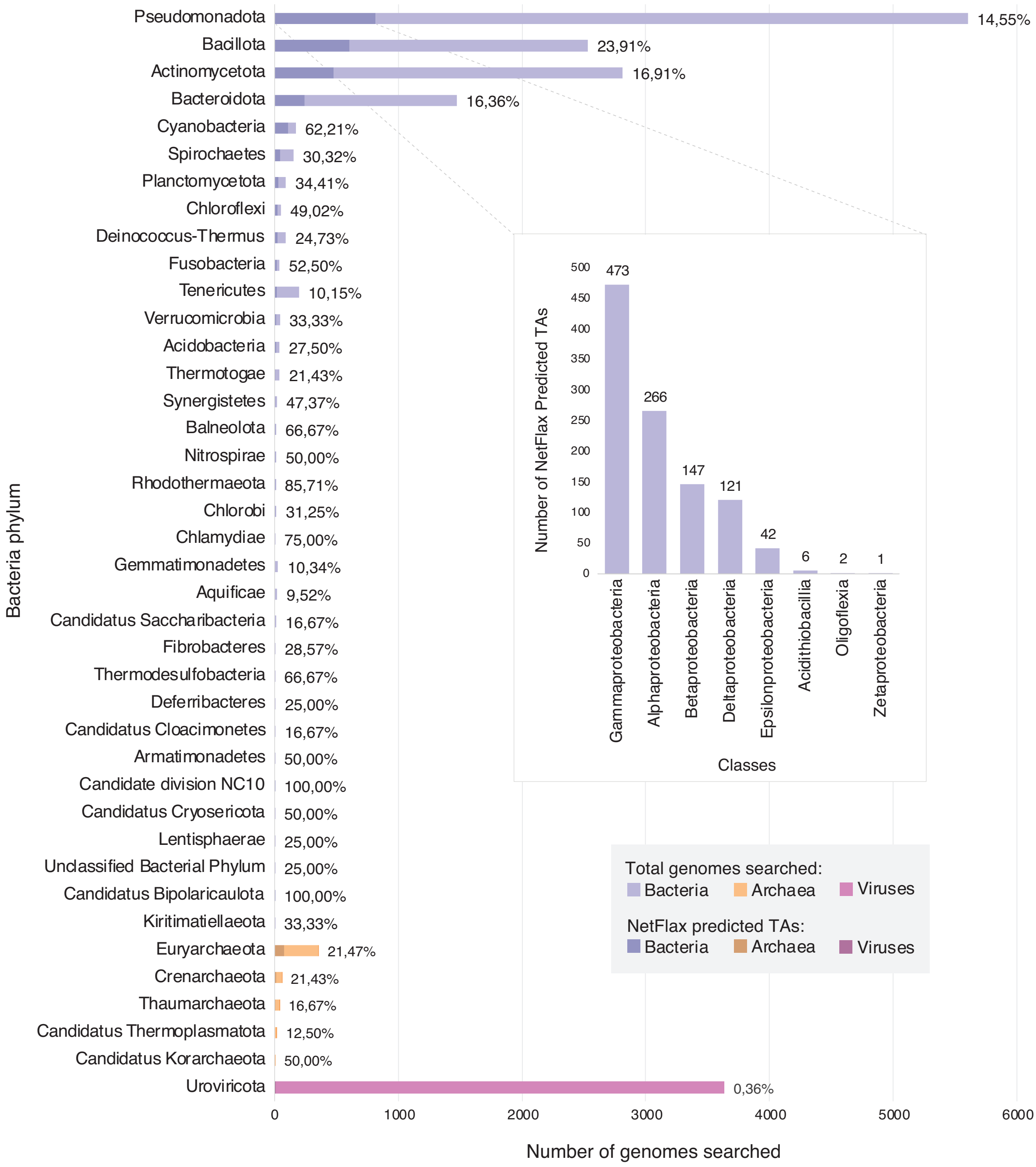


**Figure S2. The taxonomic distribution of systems in the core TA network.**
Bars represent the number of considered proteomes that we analysed (purple for bacteria, orange for archaea and pink for viruses). The darker colour on the bars represents the number of proteomes that carry NetFlax-predicted TA pairs. The percentage value denotes the proportion of all proteomes in each super kingdom that have predicted TAs. Inset: the number of predicted TAs in different Pseudomadata (Proteobacteria) taxonomic classes.

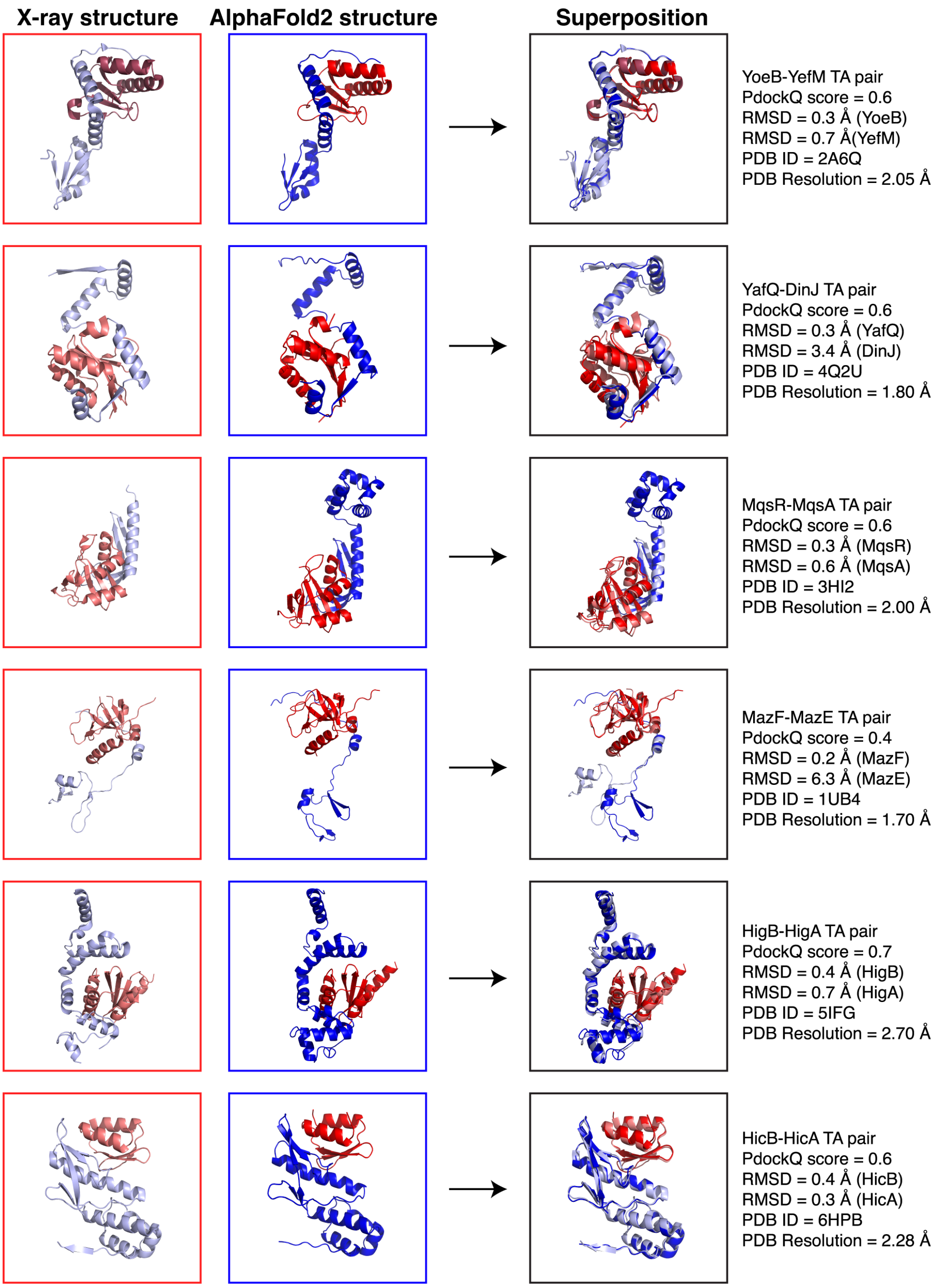


**Figure S3. A comparative depiction and analysis of the AlphaFold-generated TA pair complexes (central panel) against their respective crystal structures (left panel).**
In this analysis the following TA pairs were considered YoeB-YefM (PDB: 2A6Q), YafQ-DinJ (PDB: 4Q2U), RelE-RelB (PDB: 4FXE), MqsR-MqsA (PDB: 3HI2), MazF-MazE (PDB: 1UB4), HigB-HigA (5IFG) and HicB-HicA (PDB: 6HPB). The toxin structures have been highlighted in red, while the antitoxin structures have been highlighted in blue (for each TA pair). Experimentally solved TA structures are highlighted in lighter shades (for instance, light red for toxin and light blue for antitoxin) while the AlphaFold-generated models are highlighted in brighter shades. Additionally, the PDB IDs considered for the experimentally solved TA structures and their structural alignment (right panel) RMS values with the AlphaFold-generated models are also highlighted.

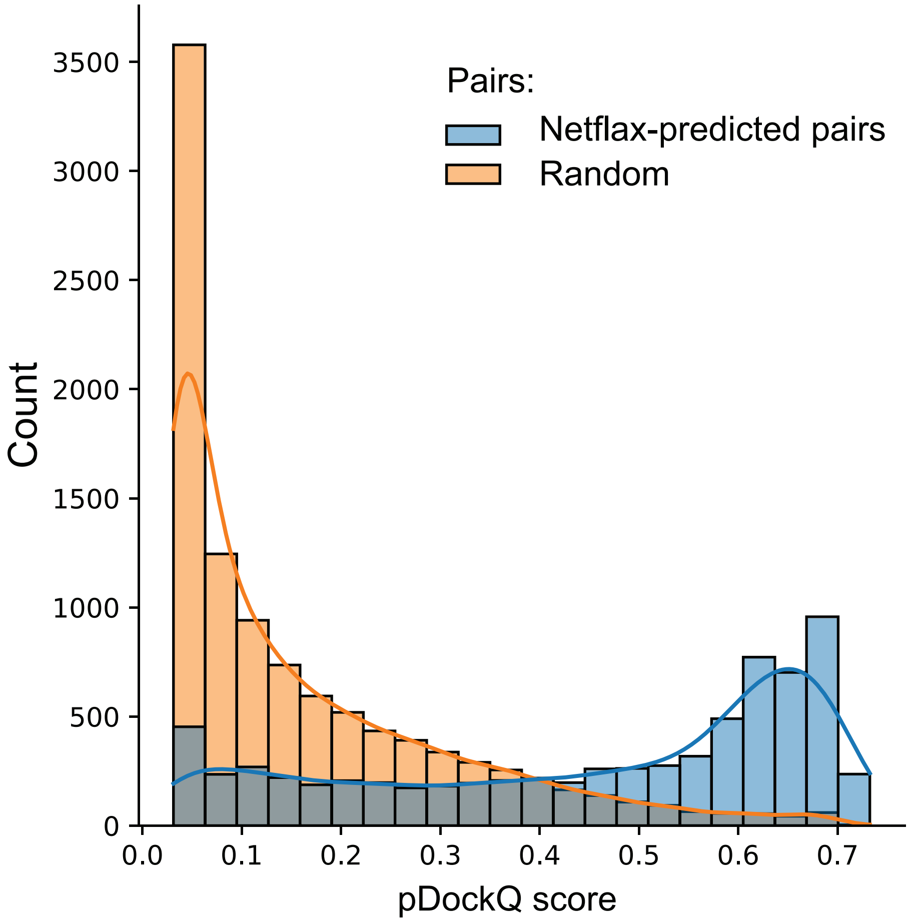


**Figure S4. Distribution of model confidence of NetFlax-predicted TA pairs (in blue) and random pairs (in orange).**
The graph is made with predicted TA pDockQ score from full (unpruned) predictions (**Dataset S1**).

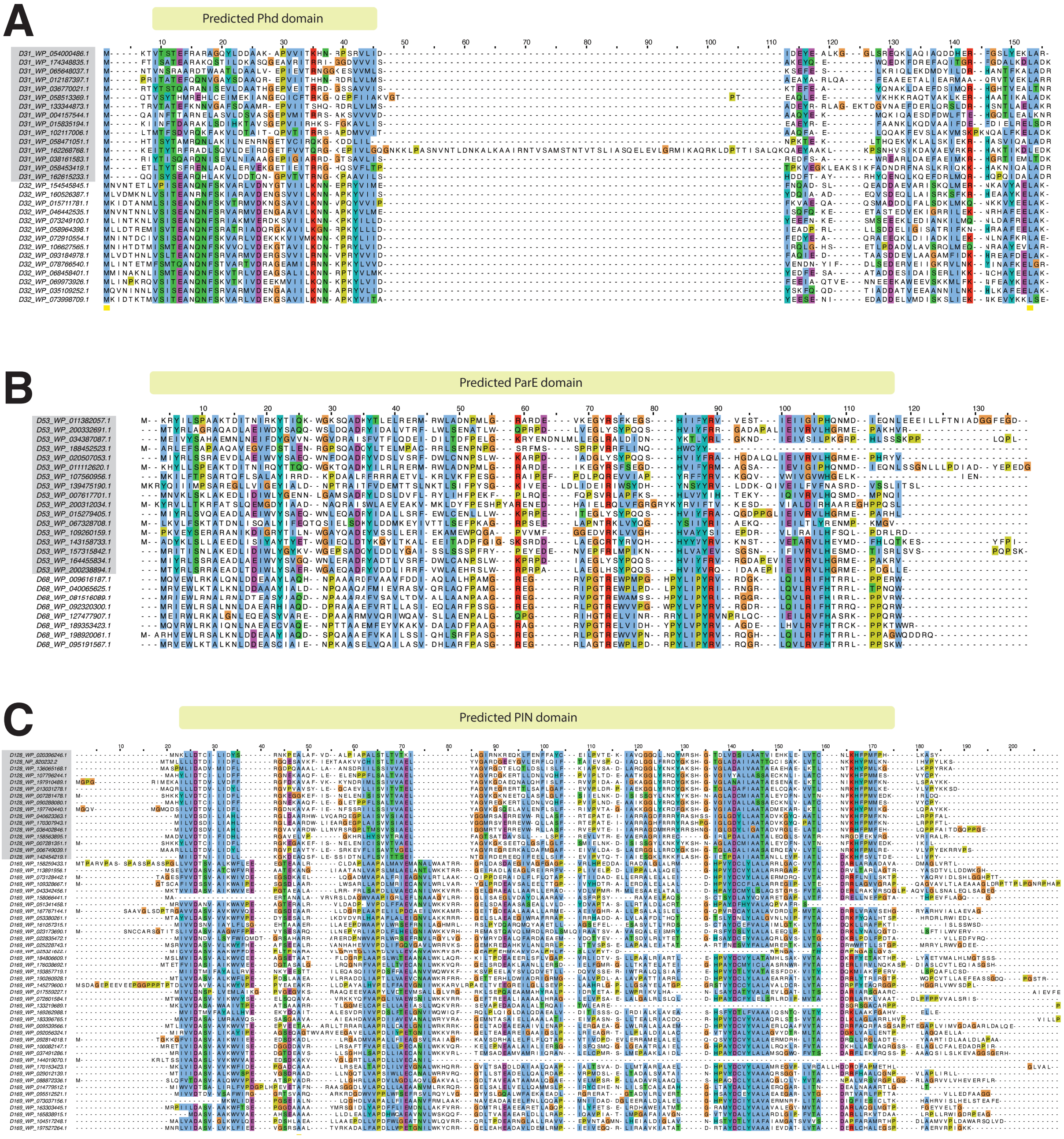


**Figure S5. Multiple sequence alignment of clusters that are related but are in different nodes.** Examples from those annotated as Phd (**A**), ParE (**B**) and PIN (**C**).

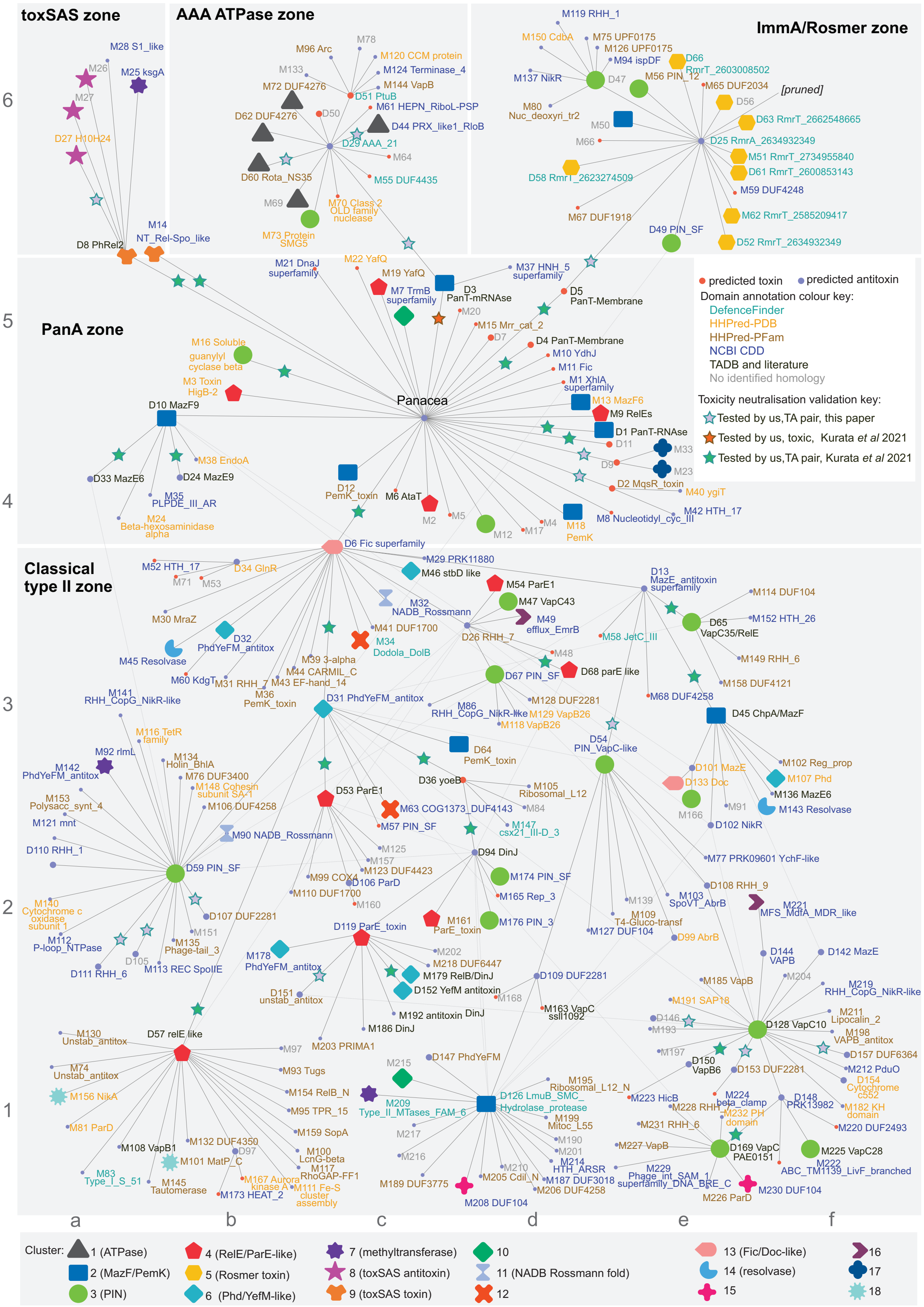


**Figure S6. Common folds mapped on to the NetFlax network.**
Symbols on nodes show the fold clusters, as per the lower legend.

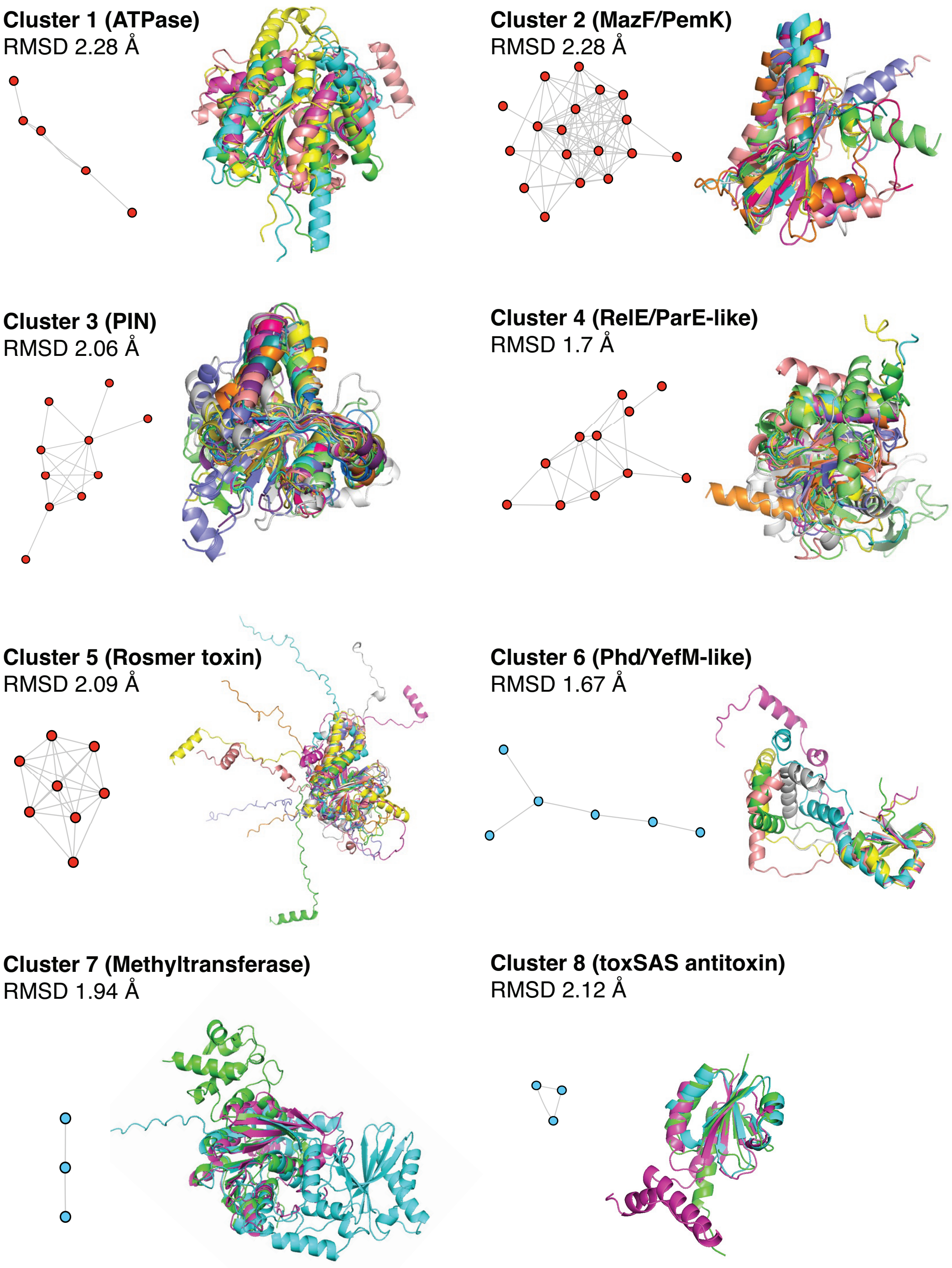


**Figure S7. Structural alignment of the major clusters of protein folds.**
The eight major clusters from our representative AF2 model dataset as aligned with the mTM-align program are shown. The RMSD (root mean squared deviation) value for each cluster is indicated. The edges of the networks are representative of the structural similarity based on an E-value below $1e^{-3}$. The network was visualized by implementing the organic layout in Cytoscape with the blue nodes representing the antitoxins and red nodes representing the toxin.

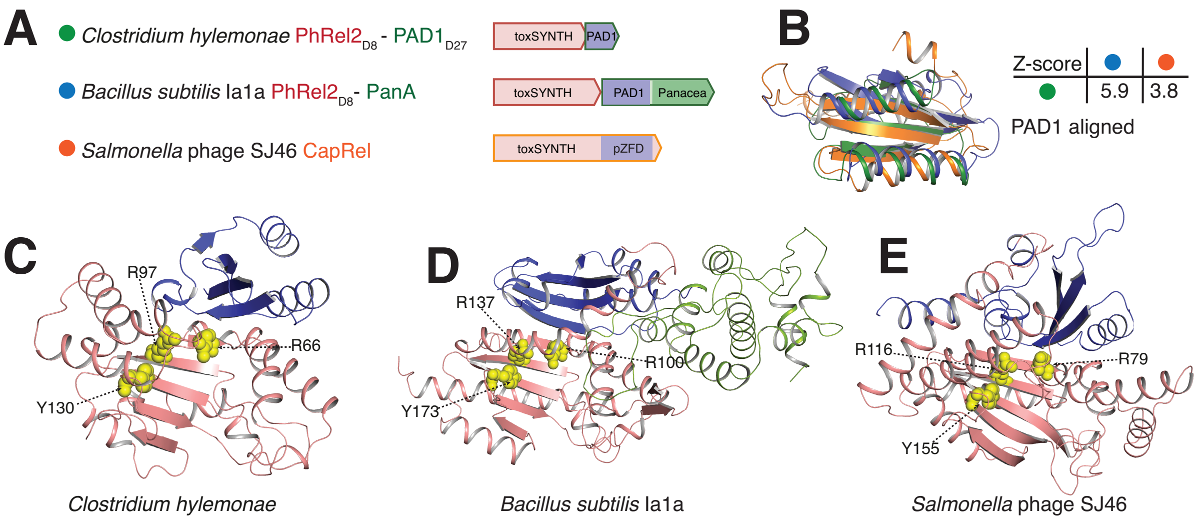


**Figure S8. Structural alignment of toxSASs and their antitoxins**.
(**A**) The reading frame and domain boundaries of toxSAS systems.
(**B**) PAD1 domains aligned to each other with the respective Z-score indicated. PAD1 domain (blue) is present in (**C**) AlphaFold-generated models of toxSAS toxin-antitoxin complexes from *C. hylemonae* DSM 15053 PhRel2$_{D8}$–PAD1$_{D27}$ and (**D**) *B. subtilis* la1a PhRel2$_{D8}$–PanA. (**E**) The fused CapRel toxSAS from *Salmonella* phage SJ46 has an antitoxin region with a similar fold to PAD1. The active center of the toxSYNTH domain is highlighted with yellow spheres.

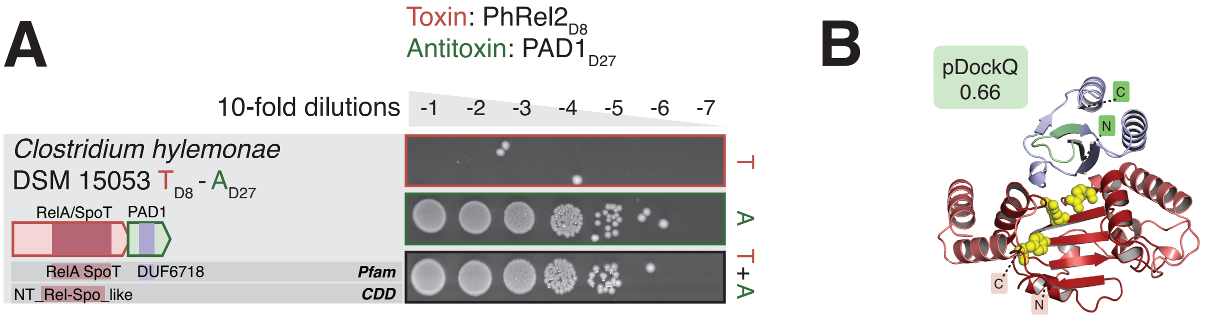


**Figure S9. Direct neutralisation of the *Clostridium hylemonae* DSM 15053 toxSAS PhRel2$_{D2}$ toxin by the PanA-associated domain 1 (PAD1$_{D28}$) antitoxin.**
(**A**) Domain organisation and TA validation though toxicity neutralisation assay.
(**B**) AlphaFold-generated structural model. The active centre of the toxSYNTH domain is highlighted with yellow spheres.

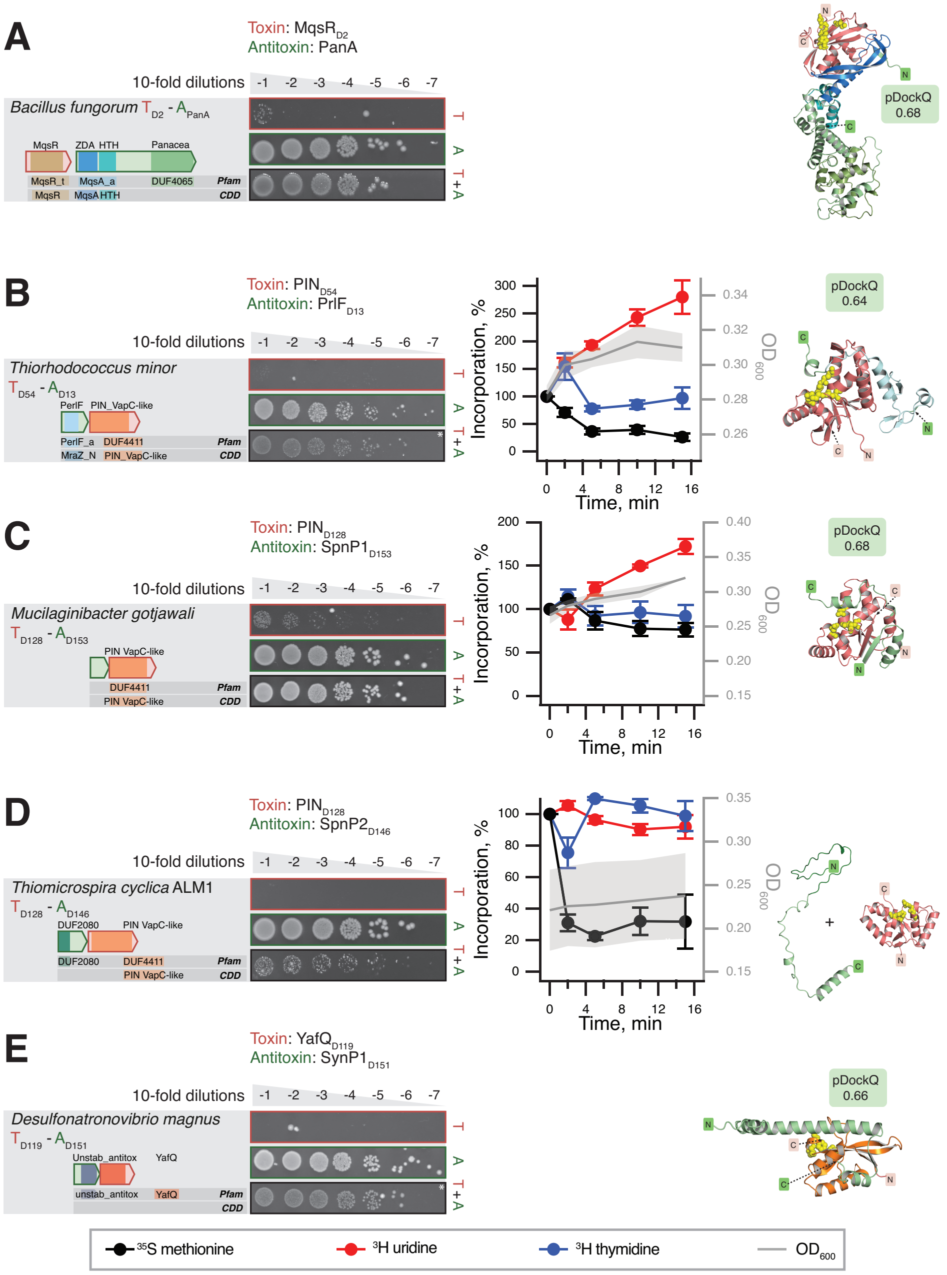


**Figure S10. TA systems containing diverse nuclease effectors.**
Domain organisation and TA validation though toxicity neutralisation assays (*left*), metabolic labelling assays with toxins expressed in wild-type *E. coli* BW25113 (*center*) and AlphaFold-generated structural models (*right*) for TAs with diverse nuclease effectors: (**A**) *B. fungorum* MqsR$_{D2}$:PanA, (**B**) *T. minor* PIN$_{D54}$:PrlF$_{D13}$, (**C**) *M. gotjawali* PIN$_{D128}$:SpnP1$_{D153}$, (**D**) *T. cyclica* AML1 PIN$_{D128}$:SpnP2$_{D146}$, and (**E**) *D. magnus* YafQ$_{D119}$:SynP1$_{D151}$. The active centre of nucleases is indicated with yellow spheres.

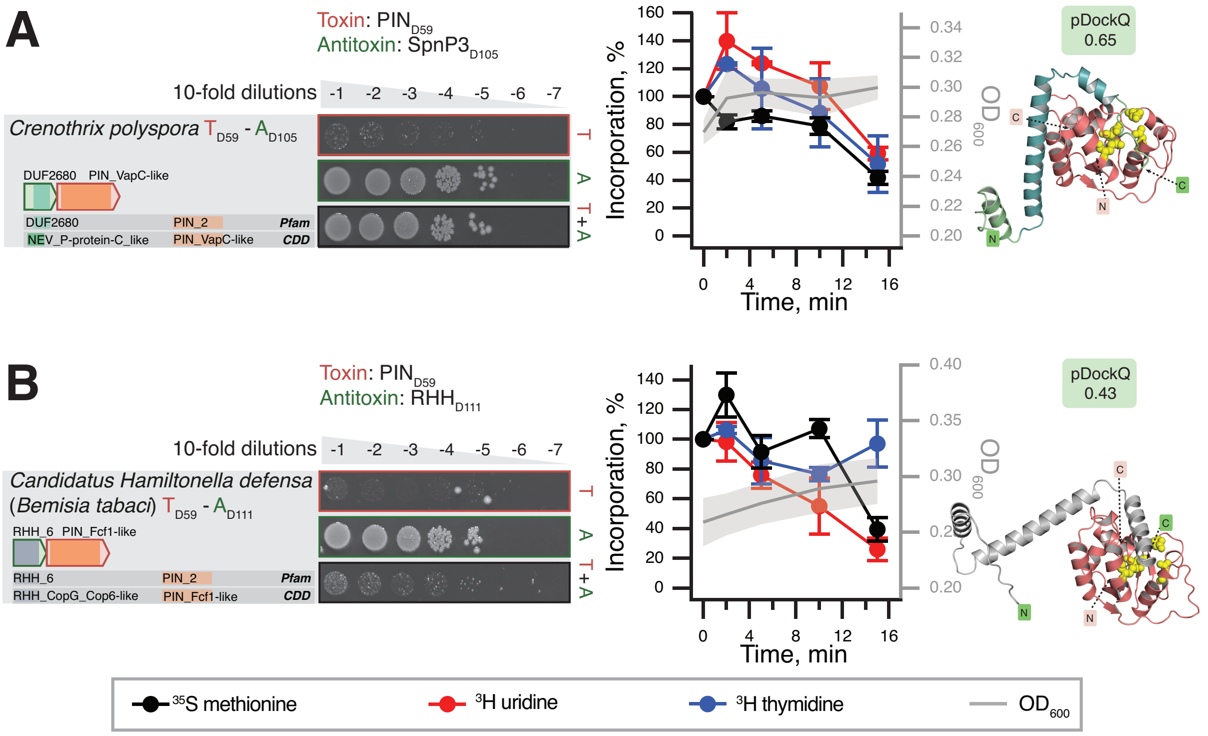


**Figure S11. TA systems containing PIN$_{D59}$ nuclease toxins.**
Domain organisation and TA validation though toxicity neutralisation assays (*left*), metabolic labelling assays with toxins expressed in wild-type *E. coli* BW25113 (*center*) and AlphaFold-generated structural models (*right*) for TAs with nuclease effectors for (**A**) *C. polyspora* PIN$_{D59}$:SpnP3$_{D105}$ and (**B**) *C. Hamiltonella defensa* PIN$_{D59}$:RHH$_{D111}$. The location of the PIN toxin active centre is highlighted with yellow spheres.

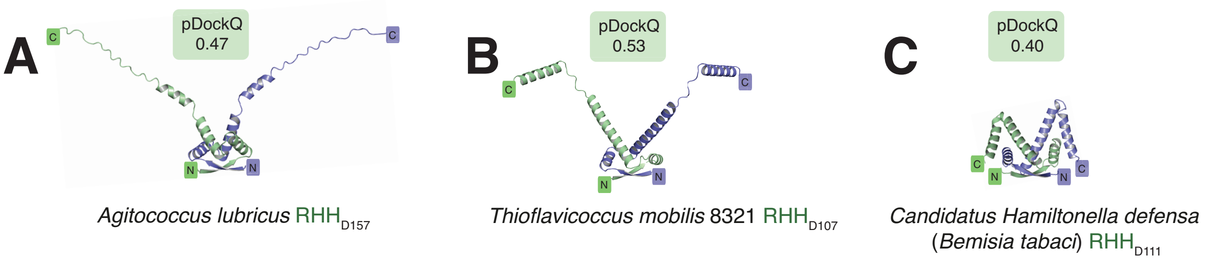


**Figure S12. AlphaFold2-predicted dimers of RHH domain-containing antitoxins.**
